# Function of FMRP domains in regulating distinct roles of neuronal protein synthesis

**DOI:** 10.1101/2021.11.15.468563

**Authors:** Michelle Ninochka D’Souza, Sarayu Ramakrishna, Bindushree K Radhakrishna, Vishwaja Jhaveri, Sreenath Ravindran, Lahari Yeramala, Dasaradhi Palakodeti, Ravi S Muddashetty

## Abstract

The Fragile X Mental Retardation Protein (FMRP) is an RNA Binding Protein that regulates translation of mRNAs, essential for synaptic development and plasticity. FMRP interacts with a specific set of mRNAs and aids in their microtubule dependent transport and regulates their translation through its association with ribosomes. However, the biochemical role of individual domains of FMRP in forming neuronal granules and associating with microtubules and ribosomes is currently undefined. Here, we report that the C-terminus domain of FMRP is sufficient to bind to ribosomes as well as polysomes akin to the full-length protein. Furthermore, the C-terminus domain alone is essential and responsible for FMRP-mediated translation repression in neurons. However, FMRP-mediated puncta formation and microtubule association is favored by the synergistic combination of FMRP domains and not by individual domains. Interestingly, we show that the phosphorylation of hFMRP at Serine-500 is important in modulating the dynamics of translation by controlling ribosome/polysome association. This is a fundamental mechanism governing the size and number of FMRP puncta, which appear to contain actively translating ribosomes. Finally through the use of pathogenic mutations, we emphasize the hierarchy of the domains of FMRP in their contribution to translation regulation.

## Introduction

Fragile X Syndrome (FXS) is a trinucleotide repeat expansion disorder of the *FMR1* gene resulting in intellectual disability (1). The silencing of the gene due to the expansion of CGG repeat in its 5’UTR results in the loss of the encoded protein Fragile X Mental Retardation Protein (FMRP). FMRP is an RNA binding protein, structurally comprising of protein interacting Tudor domains and three RNA binding domains: two K homology domains and one RGG domain in the intrinsically disordered C-terminus(2,3). FMRP is implicated in a range of biological processes such as chromatin remodeling, ion channel stability, RNA transport, ribosome heterogeneity and translation regulation (4–10). However, the contribution of FMRP to FXS pathology is mostly studied in the absence of the protein and the role of the individual domains of FMRP in these cellular processes is yet to be investigated.

FMRP plays a key role in maintaining synaptic development and plasticity through regulation of protein synthesis (11,12). FMRP is primarily shown to act as a translational repressor (13). However, FMRP also activates the translation of specific candidates both on both mGluR and NMDAR stimulation (8,9). It is important to note that FMRP bound targets localize to neuronal processes in the form of granules in a microtubule-dependent manner (14) and the expression of these targets is guided through FMRP’s distribution to polysomes (15–17). The interaction of FMRP with ribosomes seems to be an important prerequisite for this. An added level of translation regulation lies in the phosphorylation of FMRP. Phosphorylation of FMRP, primarily at Serine-500, is shown to influence the bidirectional role of FMRP as a regulator of synaptic protein synthesis(18–21).

The function of FMRP comprises the multistep process of granule formation, association with microtubule and interaction with ribosomes. Only a few studies address the mechanism of these interactions through truncations and mutations of FMRP. The N-terminus domain of FMRP is shown to modulate the expression and activity of several ion channels (22,23) while the RGG domain in the C-terminus is implicated in RNA binding and consequent gene expression (24,25). Further, the *I304N* mutation of FMRP, which results in a severe form of FXS, illustrates the importance of KH domains in polysome association and translation regulation (16,26). Apart from *I304N*, gene sequencing studies have uncovered a series of other mutations in *FMR1* leading to FXS-like phenotypes. *In-silico* analysis of *FMR1* SNPs indicated that *R138Q* mutation in the NLS of FMRP alters the localization of FMRP without any structural effect on the protein(2,27). But, mutations in the KH domains like *G266E* and *I304N* seem to disturb the interactions between domains, disrupting the overall structure and function of FMRP (27,28). C-terminus domain mutations such as *R534H* and *G482S* have been predicted to alter RNA binding and recognition (29). In addition, the frame shift mutation (*G538fs*23*) in the open reading frame of FMRP has been shown to introduce novel sequences, which drastically alter the cellular localization of FMRP (30). These mutational studies emphasize the probable structural defects and the contribution of individual domains of FMRP. However, the consequence of these point mutations in cellular processes is yet to be addressed. Further, this has generated a vacuum in the information available to understand the functions manifested by FMRP mutants.

The function of FMRP is complex and studying the role of individual domains and their synergistic effect is beneficial in understanding FMRP-mediated regulation of protein synthesis. Here, we systematically investigate the contribution of FMRP and its domains in the dynamic processes of (i) ribosome binding ii) mRNP puncta formation, and (iii) microtubule association. Our results provide structural insights on the importance of the C-terminal domain in ribosome binding and subsequent regulation of neuronal translation. We also emphasize the role of phosphorylation of FMRP in the formation of puncta and their association with microtubules. Our findings elucidate the functional consequences of pathogenic point mutations on regulatory mechanisms of FMRP. This is important to understand FMRP domains and their role in coordinating the mechanisms leading to translation regulation.

## Results

### C-terminus of FMRP is responsible for direct binding to ribosomes

FMRP is a modulator of activity-mediated translation in the nervous system (31–33) and an important requirement for this function is the ability of FMRP to interact with ribosomes and polysomes (12,33). This feature was clearly shown in I304N FMRP mutant, which does not associate with polysomes, but retains its RNA binding capacity(16). FMRP is a multi-domain protein and it is important to understand which of these domains is essential for its interaction with ribosomes. We fragmented human FMRP into three parts, the N-terminus containing Tudor domains (N-term), the KH1 and KH2 domains (collectively called the KH domain here) and the C-terminus containing the RGG box. To address the contribution of these domains of FMRP in ribosome association, first we used HEK293T cell lysate. Recombinant proteins corresponding to N-term (1-206 aa), KH domains (207-422 aa) and C-term (423-632 aa) of FMRP (Fig 1A) bearing N-terminal GST tag were expressed in *E*.*coli* Rosetta DE3 cells. We used a Baculoviral based SF9 insect cell system to express 6His-tagged full-length human FMRP (His-FMRP) since its expression in *E*.*coli* resulted in multiple truncated versions of the protein (**Fig 1A**).

**Figure 1:**
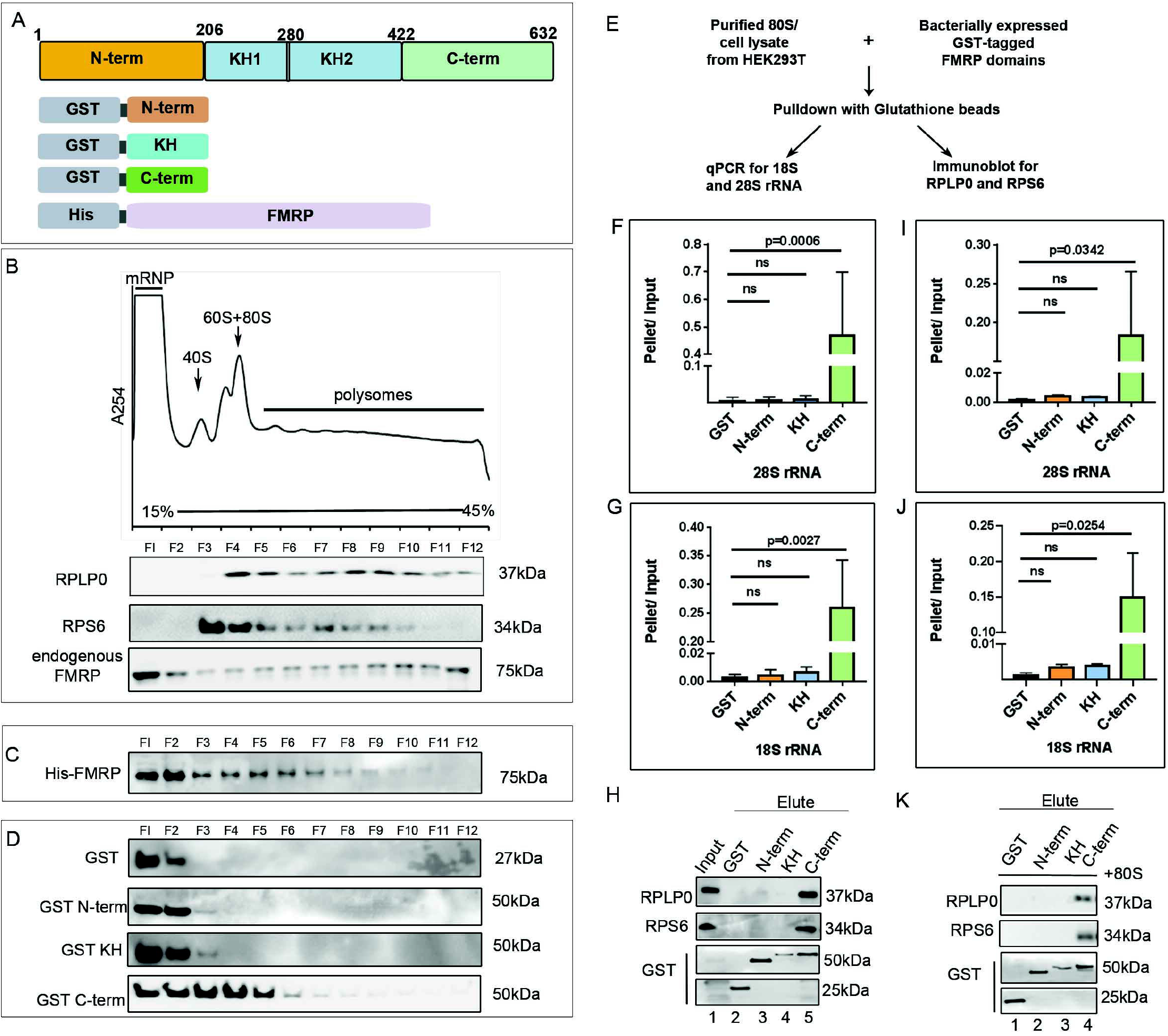
C-terminus of FMRP is important for binding to ribosomes. A. Schematic depiction of the constructs of the purified proteins for individual domains and full length human FMRP. The domains were GST tagged and expressed in bacteria. Full length FMRP was His tagged and expressed in Sf9 insect cells. The purified domains and full length FMRP was spiked into HEK293T cell lysate and subjected to polysome profiling on a linear sucrose gradient. B. Top - Representative polysome trace of spiked HEK293T cell lysate on 15%-45% linear sucrose gradient. Bottom – Representative immunoblots for endogenous FMRP, RPLP0 and RPS6 distribution in the 12 fractions obtained from polysome profiling. (n=3). C. Representative immunoblot for spiked His-tagged full length FMRP distribution in the 12 fractions obtained from polysome profiling probed with anti-His antibody (n=3). D. Representative immunoblots indicating the distribution of spiked GST-tagged FMRP domains and purified GST in the 12 fractions obtained from polysome profiling. Blots were probed with anti-GST antibody. (n=3) E. Experimental workflow to validate the binding of FMRP domains to ribosomes. F. qPCR quantification of 28SrRNA from HEK293T cell lysate bound to GST tagged FMRP domains. Data represented as mean +/-SEM, One-Way ANOVA p=0.0004 followed by Dunnett’s multiple comparisons test. G. qPCR quantification of 18SrRNA from HEK293T cell lysate bound to GST tagged FMRP domains. Data represented as mean +/-SEM, One-Way ANOVA p=0.0020 followed by Dunnett’s multiple comparisons test. H. Representative immunoblots for ribosomal proteins RPLP0 and RPS6 obtained from pull downs with GST-tagged FMRP domains incubated with HEK293T cell lysate (n=2). I. qPCR quantification of 28SrRNA from purified 80S ribosomes bound to GST tagged FMRP domains. Data represented as mean +/-SEM, One-Way ANOVA p=0.0332 followed by Dunnett’s multiple comparisons test. J. qPCR quantification of 18SrRNA from purified 80S ribosomes bound to GST tagged FMRP domains. Data represented as mean +/-SEM, One-Way ANOVA p=0.0244 followed by Dunnett’s multiple comparisons test. K. Representative immunoblots for ribosomal proteins RPLP0 and RPS6 obtained from pull downs with GST-tagged FMRP domains incubated with purified HEK293T 80S ribosomes (n=2).

Purified His-FMRP (150 pmoles) was spiked with the HEK293T lysate and subjected to polysome profiling. The distribution of ribosomes along the sucrose gradient was visualized through distribution of RPLP0 and RPS6 proteins, representing the large and small subunits of the ribosome respectively (**Fig 1B**). A significant part of endogenous FMRP and His-FMRP co-fractionated with both ribosomal and polysomal fractions (**Fig 1B and 1C**). To rule out a potential biochemical artifact from the spiked proteins, we also performed polysome-profiling assays by over expressing Flag-HA FMRP in HEK293T cells (**Fig S1A**). Over-expressed Flag-HA FMRP was distributed across all fractions of the linear sucrose gradient similar to the spiked His-FMRP protein (**Fig S1B**). 150 pmoles of each recombinant domain was spiked into HEK293T cell lysate followed by its separation on a 15-45% linear sucrose gradient. An antibody against the GST-tag was used for the detection of all the spiked domain proteins. N-term and KH domains were present only in the non–ribosomal fractions (fractions 1 and 2) where RPLP0 and RPS6 were absent (**Fig 1B and 1D**). This was quite unexpected since the KH domain has been previously reported to be important for polysome distribution of FMRP (16). Importantly, only the C-terminus domain could mimic the polysome distribution of the full-length FMRP protein (**Fig 1D**). Purified GST alone was spiked in HEK293T cell lysate as a control and it did not enter into either ribosomal or polysome fractions (**Fig 1D**). Thus, our results suggest that the C-terminus domain of FMRP is important for its association with ribosomes/polysomes.

To further validate the minimal domain of FMRP involved in the interaction with ribosomes, we performed pull-down assays with recombinant FMRP domains mixed with HEK293T cell lysate or purified human 80S ribosomes (**Fig 1E**). HEK293T cell lysates were spiked with 50 pmoles of GST-tagged domains and incubated with glutathione beads. Elutes from the beads were assayed for the presence of 18S and 28S rRNA by qPCR (as a readout for ribosomes) (**Fig 1E**). Our results clearly show that 18S and 28S rRNA was significantly pulled down only by C-term of FMRP (**Fig 1F and 1G**). In contrast, GST-N-term and GST-KH did not show any significant pull down of 18S and 28S rRNA, similar to control GST (**Fig 1F and 1G**). Alternatively, the elutes were also assayed for the presence of RPS6 and RPLP0 ribosomal proteins. Once again, only C-term was associated with ribosomal proteins as compared to N-term, KH and GST alone (**Fig.1H**). We performed an *in-vitro* binding assay with purified mRNA-free human 80S ribosomes and molar excess of GST-tagged domains (10X). Elutes were probed for the enrichment of 18S and 28S rRNA (**Fig 1I and J**) as well as ribosomal proteins RPLP0 and RPS6 (**Fig.1K**). Similar to the results with HEK293T lysates, we observed that only C-term of FMRP associated with purified 80S ribosomes.

### Phosphorylation of FMRP C-term determines its ribosome association

Post-translational modifications such as phosphorylation are known to modulate many functions of FMRP: RNA binding and protein-protein interactions in particular (18). The primary site for phosphorylation is Serine 500 within the C-terminus domain of hFMRP (18). Hence we postulated that altering the phosphorylation status of FMRP could alter its association with ribosomes/polysomes (6). We transfected HEK293T cells with Flag-HA tagged FMRP WT, phosphomimetic (FMRP S500D) and de-phosphomimetic (FMRP S500A) for 24h followed by separation on a linear sucrose gradient (**Fig 2A**). Under steady state conditions, FMRP WT sediments with ribosomal (fractions 3-12) as well as non-ribosomal fractions (fractions 1 and 2). Immunoblots from the same fractions, probed with an antibody against Phosphorylated Serine500 (p-FMRP), clearly showed an accumulation of phosphorylated FMRP only in the initial fractions (fractions 1-2) (**Fig 2B**). Similarly FMRP S500D was present only in the non-ribosomal fractions (fractions 1-2). FMRP S500A was distributed in all the fractions including polysomes (fractions 1-12), similar to FMRP WT (**Fig 2B**).

**Figure 2:**
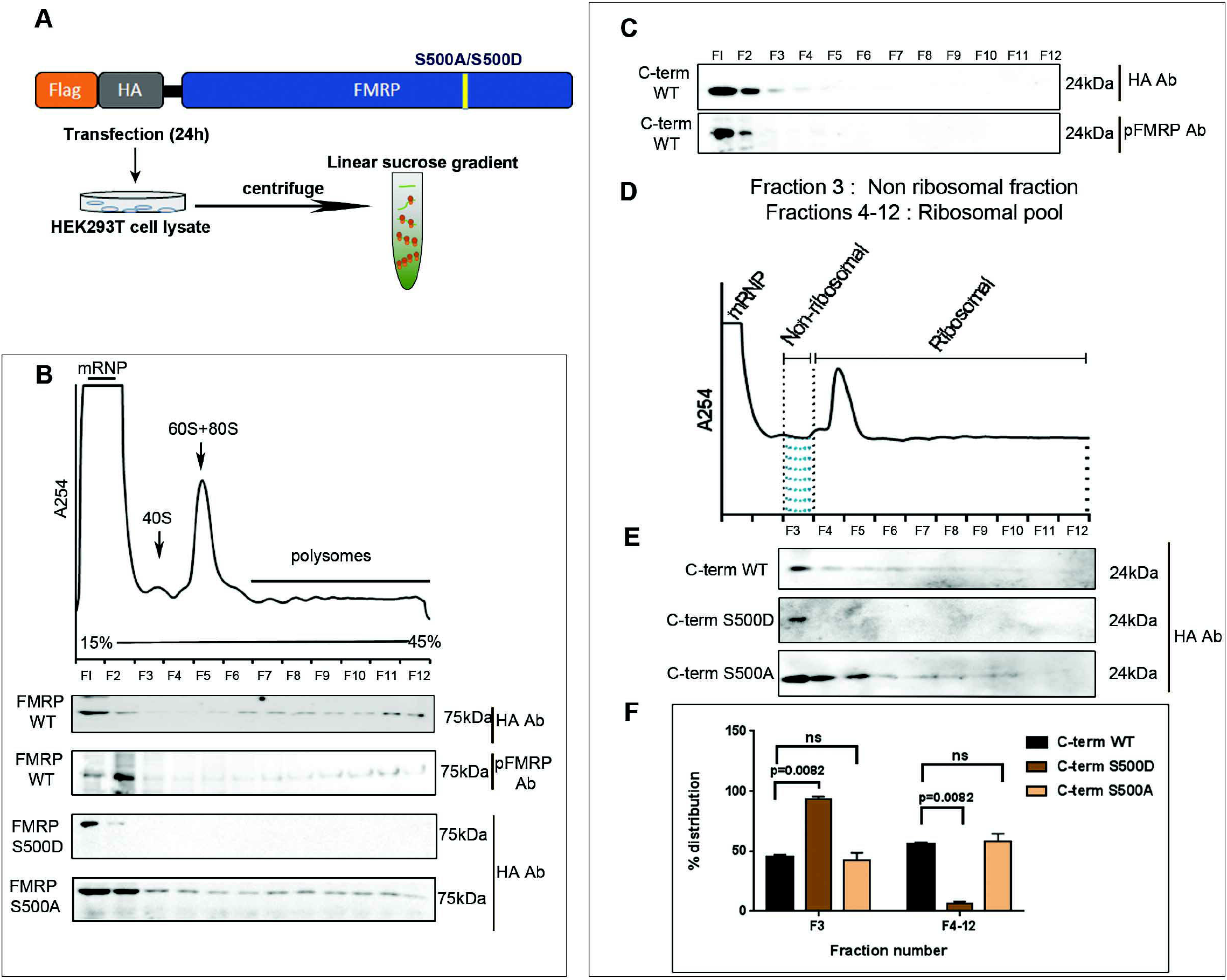
Phosphorylation of FMRP C-terminus determines its ribosome association. A. Schematic describing WT FMRP, S500D and S500A mutants of FMRP that were transfected in HEK293T cells for 24h before subjecting it to polysome profiling on a linear sucrose gradient. B. Top - Representative polysome trace obtained at 254nm indicating the division of fractions into mRNP, 40S, 60S+80S and Polysomes. Bottom – Representative immunoblots indicating the distribution of overexpressed full-length WT FMRP, phosphomimetic S500D FMRP and dephospho-mimetic S500A FMRP along the 12 fractions of linear sucrose gradient. WT FMRP transfected fractions were also probed for phospho FMRP with anti-pFMRP (Serine500) antibody (n=3). C. Representative immunoblots indicating the distribution of C-term WT overexpressed in HEK293T cells along the 12 fractions of the linear gradient. Fractions were also probed for phospho (at S500) Cterm distribution with anti-pFMRP (Serine500) antibody (n=3). D. Representative polysome profile with trace obtained at 254nm. Schematic indicates the division of fractions based on the presence of assembled ribosomal subunits. Fraction 3 corresponds to the Non-ribosomal pool. Fractions 4-12 correspond to the ribosomal pool. E. Representative immunoblots indicating the distribution of overexpressed C-term WT, phospho-mimetic (C-term S500D) and dephospho-mimetic (Cterm-S500A) along fractions 3 to 12 of a linear sucrose gradient. Blots were probed with anti-HA antibody (n=3). F. Graph indicating the quantification of overexpressed C-term variants in non-ribosomal fraction (F3) versus Ribosomal fractions (F4-12). n=3 for each condition. Data represented as mean +/-SEM, One-Way ANOVA p=0.004 for F3, p=0.004 for F4-12 followed by Bonferroni’s Multiple comparisons test.

To test this at the level of domains, we transfected HEK293T cells with Flag-HA FMRP C-term and subjected it to polysome profiling. We observed that the C-term domain, which contains the primary phosphorylation site Serine 500, was mostly restricted to the initial fractions of the sucrose gradient. Probing the same fractions with a p-FMRP antibody showed that the majority of C-term domain, which was in the initial fractions, was phosphorylated (**Fig 2C**). We investigated if altering the phosphorylation of Serine 500 in C-term could improve its distribution in the polysomal fractions. To test this, we overexpressed the phosphomimetic (C-term S500D) and de-phosphomimetic (C-term S500A) in HEK293T cells and examined their polysome distribution (**Fig 2D and 2E**). Since the overexpression of full-length FMRP, C-term and their phospho-mutants were performed in HEK293T cells in the background of endogenous FMRP; we suspected that there could be a modulation in the expression of our constructs and their association with ribosomes. Hence to understand the difference in the distribution between C-term WT, C-term S500D and C-term S500A, we quantified the amount of overexpressed protein in fraction 3 (non-ribosome) versus fractions 4-12 (ribosome+polysomes) (**Fig 2D**). C-term S500A showed a significant increase in polysome association and conversely C-term S500D showed significant accumulation in the initial fractions (**Fig 2E and 2F**). Together with our previous results, we conclude that C-term alone is sufficient to bind to the ribosome and this association is further enhanced through dephosphorylation (**Fig 2F**).

Finally, we generated a combination of KH and C-term domains (referred to as KH+C-term) and investigated its polysomal distribution (**Fig S2A**). We observed that most of the overexpressed KH+C-term was found to be in the initial fractions of the linear gradient (**Fig S2A**). Immunoblotting with p-FMRP antibody showed that accumulated KH+C-term is phosphorylated (**Fig S2A**). Similarly, the difference in ribosomal association between WT, S500D and S500A mutants of KH+Cterm was distinguishable only when quantified between fractions 3 and fractions 4-12 (**Fig S2B and S2C**). We clearly observed that the dephosphorylation of KH+Cterm significantly increased its association with the heavier ribosomal fractions (**Fig S2C**). This further confirms that the phosphorylation status of FMRP can indeed dictate its ribosome binding ability.

### The C-terminus of FMRP alone is sufficient to inhibit global protein synthesis in neurons

FMRP is primarily described as a modulator of protein synthesis in neurons, which at basal level, is shown to induce translation inhibition (34,35). But the contribution of individual domains of FMRP in regulating translation is not known. To examine this, we transfected rat primary cortical neurons with Flag-HA tagged N-term, KH and C-term of FMRP and measured *de novo* protein synthesis through FUNCAT (fluorescent non-canonical amino-acid tagging) (**Fig 3A**)(36). As expected, there was a significant reduction in the FUNCAT signal on over-expression of full-length FMRP in comparison to the untransfected neurons (**Fig 3B and 3C**). Interestingly, we found that C-term alone could significantly decrease the FUNCAT signal by 25% similar to full-length FMRP (**Fig 3B and 3C**). Although KH domains have been previously shown to be indispensable for inhibition of target mRNA translation, we did not capture any significant change in the FUNCAT signal in KH transfected neurons (**Fig 3B and 3C**). Similarly, transfection of N-term alone also did not alter the FUNCAT signal demonstrating that N-term has no effect on global translation (**Fig 3B and 3C**).

**Figure 3:**
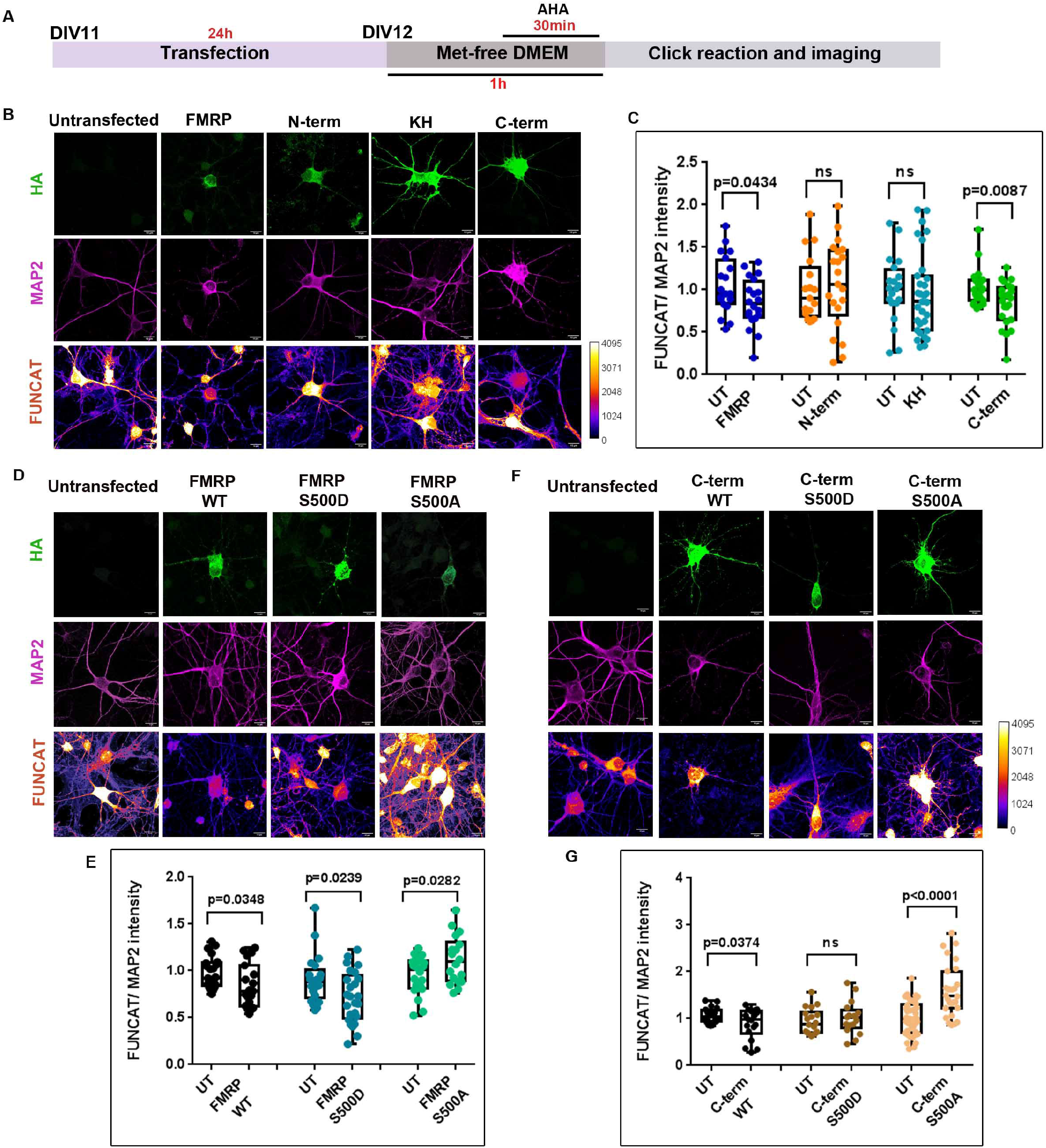
C-terminus is responsible for FMRP-mediated translation regulation in neurons. A. Experimental workflow. Primary rat cortical neurons (DIV11) were transfected for 24h with full length and FMRP domains. Transfected neurons were subjected to fluorescent non-canonical amino acid tagging (FUNCAT) along with immunostaining for HA and MAP2. B. Representative images for HA, MAP2 and FUNCAT fluorescent signal in neurons transfected with full length FMRP and FMRP domains (Scale bar - 10µm) C. Box plot representing the quantification of the FUNCAT fluorescent intensity normalized to MAP2 fluorescent intensity for full length and domains of FMRP. For each FMRP construct, the FUNCAT signal from the transfected neuron was normalized to the untransfected neuron. The box extends from 25th to 75th percentile with the middlemost line representing the median of the dataset. Whiskers range from minimum to maximum data point. Unpaired t-test for each construct, n = 15-30 neurons from 5 independent experiments D. Representative images for HA, MAP2 and FUNCAT fluorescent signal in neurons transfected with full length WT, S500D and S500A FMRP variants (Scale bar - 10µm) E. Box plot representing the quantification of the FUNCAT fluorescent intensity normalized to MAP2 fluorescent intensity for full length WT, S500D and S500A FMRP. For each FMRP construct, the FUNCAT signal from the transfected neuron was normalized to the untransfected neuron. The box extends from 25th to 75th percentile with the middlemost line representing the median of the dataset. Whiskers range from minimum to maximum data point. Unpaired t-test for each construct, n = 20-40 neurons from 4 independent experiments F. Representative images for HA, MAP2 and FUNCAT fluorescent signal in neurons transfected with C-terminus WT, S500D and S500A variants (Scale bar - 10µm) G. Box plot representing the quantification of the FUNCAT fluorescent intensity normalized to MAP2 fluorescent intensity for C-terminus WT, S500D and S500A variants. For each FMRP construct, the FUNCAT signal from the transfected neuron was normalized to the untransfected neuron. The box extends from 25th to 75th percentile with the middlemost line representing the median of the dataset. Whiskers range from minimum to maximum data point. Unpaired t-test for each construct, n = 20-40 neurons from 4 independent experiments

Previously we highlighted the role of phosphorylation in influencing FMRP-ribosome association (**Fig 2B**). We tested if the phosphorylation status of FMRP can also influence global translation regulation. We transfected rat primary cortical neurons with FMRP WT, S500D and S500A constructs for 24h prior to FUNCAT (**Fig 3D**). As indicated in earlier results, over-expression of FMRP WT significantly reduced FUNCAT signal indicating an inhibition in total protein synthesis (**Fig 3D and 3E**). Likewise, over-expression of phosphomimetic FMRP S500D also led to a reduction in the FUNCAT signal. On the contrary, FMRP S500A showed a significant increase in the FUNCAT signal in comparison to untransfected neurons (**Fig 3D and 3E**). In a parallel experiment WT, S500D and S500A mutants of C-term were transfected in neurons to determine if C-term also follows a similar mechanism in regulating translation. We captured a similar reduction in FUNCAT signal with overexpression of C-term WT and a consistent increase in FUNCAT signal with C-term S500A mutant (**Fig 3F and 3G**). However, the FUNCAT signal in the neurons transfected with C-term S500D was unchanged (**Fig 3F and 3G**). Taken together, our results conclude that the C-term of FMRP is the primary driver of translation regulation and C-term alone can capture the translation switch of FMRP by modulating its phosphorylation status.

### FMRP puncta size is regulated by phosphorylation and puncta are sensitive to Puromycin

FMRP has been shown to be a component of dynamic membrane-less granules in neurons (9,37– 39). This process of granule formation has been shown to be facilitated by phase-separation through the Intrinsically Disordered Region (IDR) of FMRP *in-vitro* (40). However, the role of other domains in formation of granules in mammalian cells is unclear. To investigate this, we over-expressed Flag-HA tagged domains of FMRP in rat primary cortical neurons and quantified the size and relative number of the neuronal puncta containing these tagged proteins (**Fig 4A and 4B**). We observed that the puncta of all individual domains of FMRP were smaller in area compared to that of full-length FMRP (**Fig 4C**). However, KH domain containing puncta were the least in number in comparison to the full-length protein (**Fig 4D**). His-GFP, which was transfected as a control, showed a diffuse pattern of expression (non punctate) compared to the punctate expression of FMRP (**Fig S3A, S3B and S3C**). Interestingly, puncta containing KH domain was not significantly different from that of GFP-containing puncta, indicating that KH domain alone has the least puncta-forming ability among other domains of FMRP (**Fig S3A, S3B and S3C**). On fusing KH to either the N-term or C-term we observed an increase in puncta size and puncta number. Although they were not as good as the full-length protein (**Fig S3D, S3E and S3F**)

**Figure 4:**
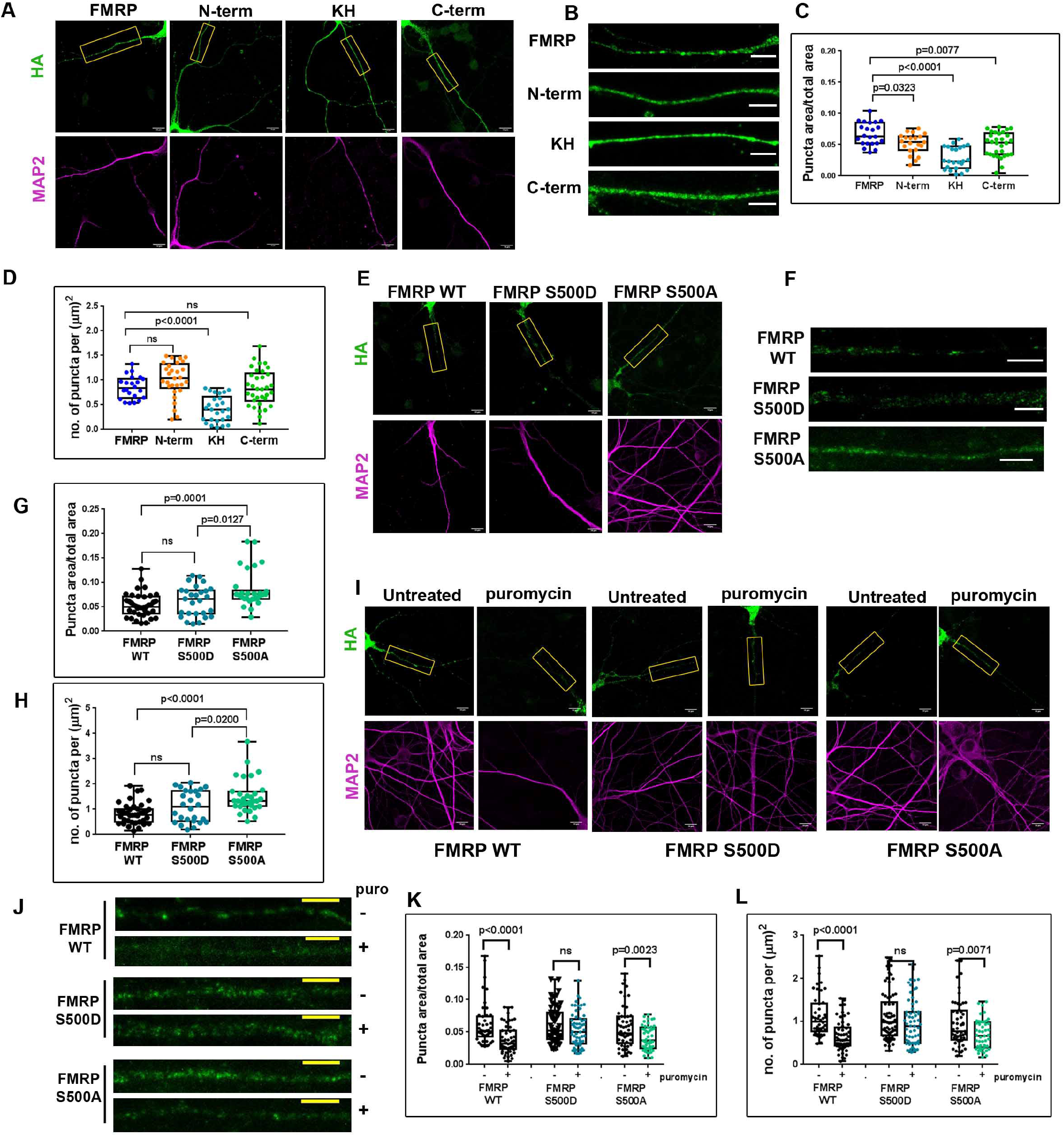
FMRP puncta size is regulated by phosphorylation and puncta are Puromycin sensitive. A. Representative images of HA and MAP2 fluorescent intensities in Primary rat cortical neurons (DIV11) transfected with full-length FMRP and FMRP domains for 24h(Scale bar - 10µm) B. Inlets shown in panel A are enlarged showing dendritic localization of puncta. (Scale bar - 5µm) C. Box plot representing the quantification of puncta area by total dendritic area for full length and domains of FMRP. The box extends from 25th to 75th percentile with the middlemost line representing the median of the dataset. Whiskers range from minimum to maximum data point. One way ANOVA p<0.001 followed by Tukey’s multiple comparison’s test n = 25-35 neurons from 5 independent experiments D. Box plot representing the quantification of the number of puncta per unit area of the dendrite for full length and domains of FMRP. The box extends from 25th to 75th percentile with the middlemost line representing the median of the dataset. Whiskers range from minimum to maximum data point. One way ANOVA p<0.001 followed by Tukey’s multiple comparison’s test, n = 25-35 neurons from 5 independent experiments E. Representative images of HA and MAP2 fluorescent intensities in Primary rat cortical neurons (DIV11) transfected with full length WT, S500D and S500A FMRP variants for 24h (Scale bar - 10µm) F. Inlets shown in panel E are enlarged showing dendritic localization of puncta. (Scale bar - 5µm) G. Box plot representing the quantification of puncta area by total dendritic area for full length WT, S500D and S500A FMRP. The box extends from 25th to 75th percentile with the middlemost line representing the median of the dataset. Whiskers range from minimum to maximum data point. One-way ANOVA p=0.0002 followed by Tukey’s multiple comparison test n = 20-40 neurons from 5 independent experiments. H. Box plot representing the quantification of the number of puncta per unit area of the dendrite for full length and WT, S500D and S500A FMRP. The box extends from 25th to 75th percentile with the middlemost line representing the median of the dataset. Whiskers range from minimum to maximum data point. One-way ANOVA p<0.0001 followed by Tukey’s multiple comparisons test. n = 25-40 neurons from 5 independent experiments. I. Representative images of HA and MAP2 fluorescent intensities in Primary rat cortical neurons (DIV11) transfected with full length WT, S500D and S500A FMRP for 24h followed by treatment with Puromycin (1mM) for 1hr (Scale bar - 10µm) J. Inlets shown in panel I are enlarged showing dendritic localization of puncta. (Scale bar - 5µm) K. Box plot representing the quantification of puncta area by total dendritic area for full length WT, S500D and S500A FMRP on Puromycin treatment. The box extends from 25th to 75th percentile with the middlemost line representing the median of the dataset. Whiskers range from minimum to maximum data point. Unpaired t-test. n = 40-60 neurons from 3 independent experiments. L. Box plot representing the quantification of the number of puncta per unit area of the dendrite for full length WT, S500D and S500A FMRP on Puromycin treatment. The box extends from 25th to 75th percentile with the middlemost line representing the median of the dataset. Whiskers range from minimum to maximum data point. Unpaired t-test. n = 40-60 neurons from 3 independent experiments

To investigate the role of phosphorylation in puncta formation, we transfected FMRP WT, FMRP S500D and FMRP S500A in rat primary cortical neurons and examined changes in puncta characteristics in neurons (**Fig 4E and 4F**). Interestingly, over-expression of FMRP S500A resulted in a significant increase in puncta area (**Fig 4G**). We also observed a significant increase in the number of puncta in neurons transfected with FMRP S500A (**Fig 4H**). FMRP-S500D containing puncta were similar in size and number to puncta containing FMPR WT (**Fig 4G and 4H**). This result indicates that dephosphorylation of FMRP has a significant influence on the size and number of FMRP containing puncta. Interestingly, this did not apply to over-expressed WT, S500D and S500A mutants of C-term and KH+C-term domains (**Fig S3G-H and S3K-L**). Altering the phosphorylation status of either C-term alone or KH+C-term did not affect the size or number of the puncta (**Fig**.**S3I-J and S3M-N**).

The molecular constituents of FMRP puncta are still unknown. It is unclear if FMRP is incorporated into translating ribosomes within these granules. To determine the translational status of FMRP containing puncta, we transfected neurons with WT, S500D and S500A FMRP and analyzed the characteristics of the puncta after treatment with protein synthesis inhibitor Puromycin (**Fig 4I and 4J**). Puromycin treatment leads to the dissociation of actively translating ribosomes/polysomes. Interestingly, on Puromycin treatment we observed a significant drop in the number and size of FMRP WT puncta (**Fig 4K and 4L**). We also observed a similar reduction in size and number of FMRP S500A puncta indicating that these puncta contain actively translating ribosomes (**Fig 4K and 4L**). S500D puncta were insensitive to Puromycin suggesting it to be consisting of stalled ribosomes (**Fig 4K and 4L**). Together, our data demonstrates that each domain of FMRP is necessary in the formation of completely functional neuronal puncta and phosphorylation modulates the characteristics of these FMRP-containing puncta.

### The KH domain of FMRP is necessary but not sufficient for FMRP-Microtubule association

The absence of FMRP-regulated local translation in response to synaptic activity is a major underlying mechanism for cognitive impairment in FXS (11,31,32). FMRP is shown to regulate the translation of a set of mRNAs in neurons through targeted localization in a microtubule-dependent manner (14). To understand this biochemical interaction of FMRP with microtubules, we employed a previously described method to stabilize and extract microtubules and its interacting molecular partners using the drug Paclitaxel/Taxol (**Fig 5A**)(33). In this assay, we observed a significant amount of endogenous FMRP present in the microtubule pellet that got further enriched on treatment with Taxol, along with tubulin (**Fig 5B**). Further on disrupting the microtubule network with Nocodazole, we observed a depletion of both tubulin and FMRP in the pellet (**Fig S4A**), confirming that the enrichment was microtubule specific. His-GFP was used as a negative control and it did not show any significant enrichment with microtubules on Taxol treatment (**Fig 5C**).

**Figure 5:**
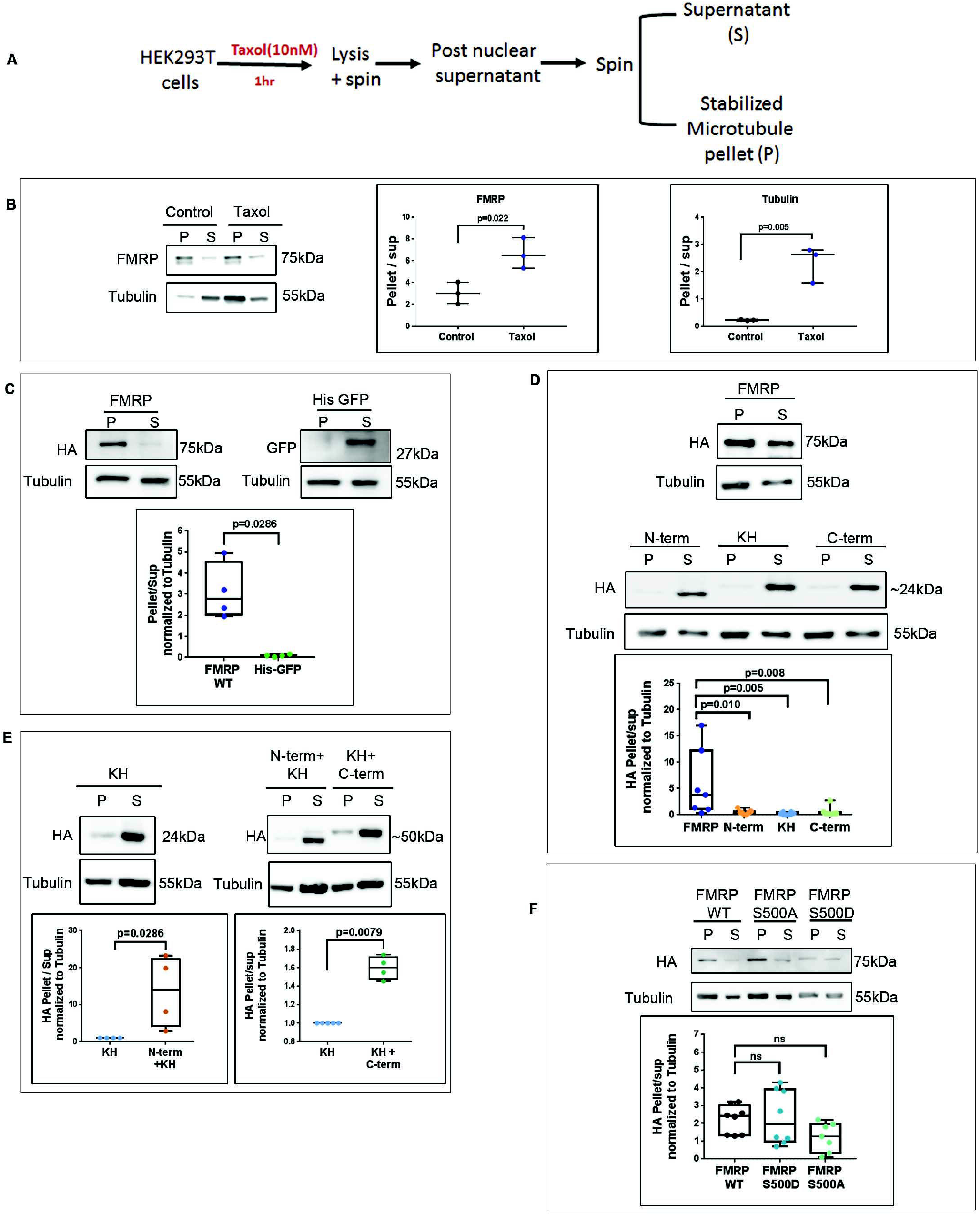
KH domain of FMRP is necessary but not sufficient enough for FMRP-Microtubule association. A.Schematic for microtubule enrichment assay on Taxol (10nM) treatment. B. Left-Representative immunoblots indicating the enrichment of endogenous FMRP and tubulin in microtubule pellet on Taxol treatment in HEK293T cells. Middle - Graph indicating the ratio of FMRP enrichment in pellet/supernatant under DMSO (control) and Taxol treated conditions. Right - Graph indicating the ratio of Tubulin enrichment in pellet/supernatant under DMSO (control) and Taxol treated conditions. Data represented as mean +/-SEM, Unpaired t-test (n=3). C. Top - Representative immunoblots indicating enrichment of WT FMRP in microtubule pellet and His-GFP in the supernatant on Taxol treatment. Bottom – box plot indicating the ratio of FMRP and His-GFP enrichment in pellet/supernatant on Taxol treatment. The box extends from 25th to 75th percentile with the middlemost line representing the median of the dataset. Whiskers range from minimum to maximum data point. Unpaired t-test (n=4). D. Immunoblots of HEK293T cell microtubule pellet enriched on Taxol treatment after transfection with full-length and FMRP domains. Box plots represent the pellet/ supernatant ratio. The box extends from 25th to 75th percentile with the middlemost line representing the median of the dataset. Whiskers range from minimum to maximum data point, One way ANOVA p=0.0053 followed by Dunnett’s multiple comparisons test (n=7-8). E. Immunoblots of HEK293T cell microtubule pellet enriched on Taxol treatment after transfection with KH, Nterm+KH and KH+Cterm. Box plots represent the pellet/ supernatant ratio. The box extends from 25th to 75th percentile with the middlemost line representing the median of the dataset. Whiskers range from minimum to maximum data point. Unpaired t-test (n=4-5). F. Immunoblots of HEK293T cell microtubule pellet enriched on Taxol treatment after transfection with WT, S500D FMRP and S500A FMRP. Box plots represent the pellet/ supernatant ratio. The box extends from 25th to 75th percentile with the middlemost line representing the median of the dataset. Whiskers range from minimum to maximum data point, One way ANOVA p=0.1325 followed by Dunnett’s multiple comparisons test (n=7-8).

Previously it has been shown that the interaction of FMRP with microtubules is RNA-dependent (33). To investigate this, we treated HEK293T cell lysate with RNase and observed a loss of FMRP from the microtubule-enriched pellet, confirming that FMRP-microtubule interaction is RNA-dependent (**Fig S4B**). To dissect out the contribution of the domains of FMRP in this interaction, HEK293T cells were transfected with N-term, KH and C-term of FMRP followed by microtubule enrichment. Surprisingly, microtubule binding of all the individual domains of FMRP was significantly lower than that of the full-length FMRP protein (**Fig 5D**). Next, we used N-term+KH and KH+C-term constructs in our microtubule-enrichment assay and observed that the combinations of domains significantly improved microtubule binding (**Fig 5E**). The role of phosphorylation in FMRP-microtubule association has been poorly studied. To investigate this, we performed microtubule-enrichment assays with HEK293T cells transfected with FMRP WT, S500D and S500A constructs. Interestingly, neither of the phosphomutants showed any significant difference in comparison to WT with respect to microtubule association implying that this post-translational modification does not influence microtubule binding (**Fig 5F**).

### Single nucleotide mutations from patients reveal functional hierarchy in FMRP domains

Fragile X Syndrome (FXS) results from the silencing of the *FMR1* gene due to the expansion of an unstable triplet CGG motif present in the 5’UTR (34). Nonetheless, high throughput sequencing studies have also revealed the presence of multiple single point mutations that render FMRP non-functional (28,41,42). We made use of relevant pathogenic mutations in the N-term (*R138Q*), KH domains (*I304N, G266E*) and C-term (*G482S, R534H, G538fs*23*) of FMRP to understand the contribution of individual domains in its function (27,29,30). We sought to investigate the impact of these mutations on the role of FMRP as a translation regulator. We generated Flag-HA FMRP constructs bearing selected single point mutations in their respective domains (**Fig 6A**). Additionally, we also generated a functionally relevant mutation in the KH1 domain (*I241N*) to mimic the well-studied patient identified mutation *I304N* that is present in the KH2 domain (**Fig 6A**) (15). First, we tested the impact of these mutations in ribosome association through linear sucrose density centrifugation. We observed that *I241N, I304N* and *G266E* mutations, residing in the KH domains of FMRP, drastically reduced its association with ribosomes and heavy polysomes presumably by affecting the RNA binding capacity of the protein (**Fig 6B**). *R138Q* mutation in FMRP structurally preserves all the RNA binding domains and hence did not show any significant difference in polysome distribution in comparison to the WT protein as previously described (**Fig 6B**)(2). Similarly the C-term mutations *R534H* and *G482S* showed no change in polysome distribution in comparison to FMRP WT (**Fig 6B**). *G538fs*23* mutation disrupts the RGG domain and generates an additional 23 novel amino acids in the C-term. This alteration of the C-term resulted in loss of ribosome/polysome binding (**Fig 6B**).

**Figure 6:**
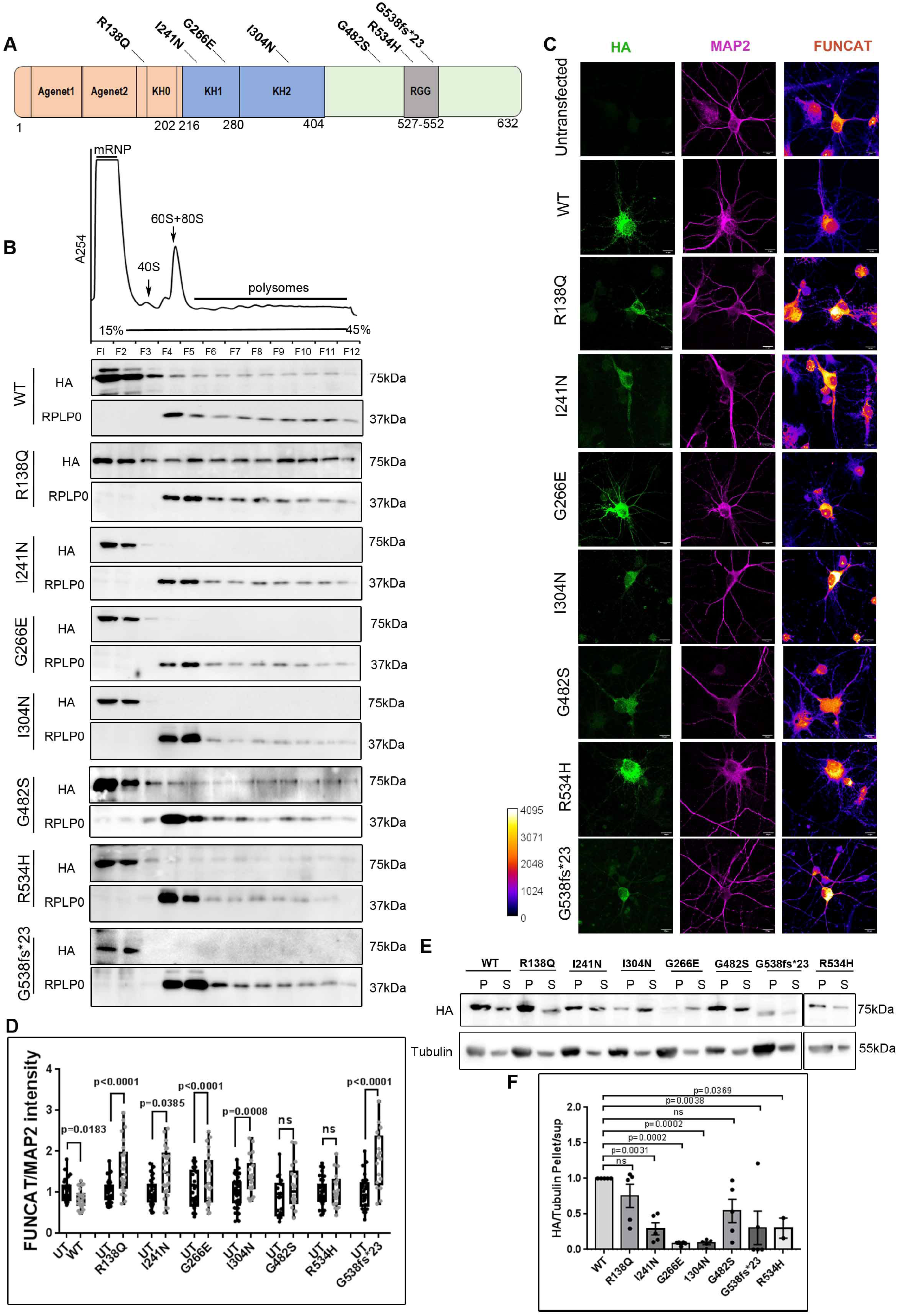
KH domain mutations drastically alter FMRP’s association with ribosomes and microtubules. A. Schematic depicting position of mutations in FMRP identified in patients with FXS and intellectual disability. B. Top - Representative polysome trace of HEK293T cell lysate transfected with full length WT and mutants for 24h. Bottom – Representative immunoblots indicating the distribution of FMRP mutants (probed with HA antibody) with corresponding distribution of ribosome fractions (probed with RPLP0 antibody). C. Representative images for HA, MAP2 and FUNCAT fluorescent signal in neurons transfected with full length WT and FMRP mutants (Scale bar - 10µM) D. Box plot representing the quantification of the FUNCAT fluorescent intensity normalized to MAP2 fluorescent intensity for WT and mutants of FMRP. For each FMRP construct, the FUNCAT signal from the transfected neuron was normalized to the untransfected neuron. The box extends from 25th to 75th percentile with the middlemost line representing the median of the dataset. Whiskers range from minimum to maximum data point. Unpaired t-test, n= 20-35 neurons from 6 independent experiments. E. Representative immunoblots indicating the enrichment of over-expressed WT FMRP, FMRP mutants and tubulin in microtubule pellet and supernatant on Taxol treatment in HEK293T cells (n=3-5). F. Box plots indicating the ratio of enrichment of WT and FMRP mutants in pellet/supernatant on Taxol treatment of HEK293T. The box extends from 25th to 75th percentile with the middlemost line representing the median of the dataset. Whiskers range from minimum to maximum data point, One way ANOVA p=0.0002, followed by Dunnett’s multiple comparisons test (n=3-5).

Next we sought to validate the functional effect of these mutations on translation regulation. Rat primary cortical neurons were transfected with Flag-HA FMRP mutants for 24h and changes in global protein synthesis was assayed using FUNCAT. As shown previously, FMRP WT transfected neurons showed an overall reduction in the FUNCAT signal (**Fig 6C and 6D**). This is in agreement with the general role of FMRP as an inhibitor of translation (**Fig 3B-E**). On the contrary, while the KH domain mutations *I241N, G266E* and *I304N* failed to inhibit translation, they also caused a significant increase in FUNCAT signal in comparison to untransfected neurons (**Fig 6C and 6D**). Neurons transfected with the C-term mutants *G482S* and *R534H* did not exhibit any change in overall FUNCAT signal in comparison to respective untransfected neurons indicating that these mutations abolish the translation inhibitory function of FMRP (**Fig 6C and 6D**). The C-term *G538fs*23* mutant also showed an increase in FUNCAT signal in the respective neurons (**Fig 6C and 6D**). This gain-of-function could be attributed to the insertion of novel amino acid sequences within the C-term. Although our initial data indicated that *R138Q* mutation does not affect ribosome binding, we observed an increase in FUNCAT signal in comparison to untransfected neurons (**Fig 6C and D**). Hence we postulate that mutations might alter the characteristics of their respective domains thereby resulting in differential regulation of global translation.

Further, we sought to examine the effect of these mutations on microtubule binding. We transfected HEK293T cells with Flag-HA FMRP mutants and followed it by a microtubule enrichment assay. KH domain mutations such as *I241N, G266E* and *I304N* did not get significantly enriched in the microtubule pellet in comparison to FMRP WT (**Fig 6E and 6F**). Similarly C-term mutations *R534H* and *G538fs*23* also significantly hampered FMRP’s association with microtubules compared to FMRP WT (**Fig 6E and 6F**). However, we did not observe any significant effect of *R138Q* and *G482S* on FMRP-microtubule association (**Fig 6E and 6F**). Together, our data concludes that distinct domains contribute to the precise functioning of FMRP in processes leading to translation regulation. Further, pathogenic mutations identified in the individual domains of FMRP affect their ability to associate with and regulate various components of translation.

## Discussion

Here we investigate the contribution of individual domains of FMRP in its primary role of translation regulation. On dissecting FMRP into single domains and domain combinations, we elucidate the independent and collective roles of the N-terminus, KH and C-terminus domains in the processes of granule formation, microtubule binding and ribosome association. Our in-vitro experiments and overexpression studies led us to an important conclusion that the C-terminus domain of FMRP is extremely essential for its binding to the ribosome. To this extent we also observe that the C-terminus domain alone is sufficient to inhibit protein synthesis in a manner similar to that of the full-length FMRP. An important feature of our study is that we were able to capture the dual role of FMRP as an activator and inhibitor of protein synthesis and this was primarily guided by its phosphorylation status. We show that phosphorylation is a principal post-translational modification (PTM) of FMRP that acts as a molecular switch to regulate the majority of its functions. Evidently there also exists a portion of FMRP that is not associated with translating polysomes. Our study also focuses on the alternate aspect of translation regulation whereby FMRP gets sequestered into mRNP containing puncta that associate with microtubules. Unlike ribosome binding, our results reveal that single domains of FMRP cannot sufficiently mediate puncta formation and microtubule association. But in fact it requires the synergistic contribution from multiple individual domains to perform these functions efficiently. FMRP has been proposed to regulate the translation of a large set of dendritic mRNAs (6,35,36). Alternatively, FMRP is also known to globally inhibit protein synthesis in neurons (26). The molecular mechanism underlying FMRP-mediated translation of these subsets of mRNA has been thoroughly studied (1,6). But we focused on understanding the mechanism of FMRP-mediated translation independent of sequence or structural features of the mRNA. It also directed our attention to determine which of the domains of FMRP are implicated in this process. Previously it was shown that hFMRP inhibits translation through direct binding of the C-terminus RGG domain to the ribosome (24). Likewise, our *in-vitro* binding assays indicate that the C-terminus domain alone can directly bind to the 80S ribosome while the N-terminus and KH domains cannot (**Fig.7A**). A critical finding was that the C-terminus domain alone was sufficient enough to mimic ribosomal and polysomal association of the full-length FMRP.

**Figure7:**
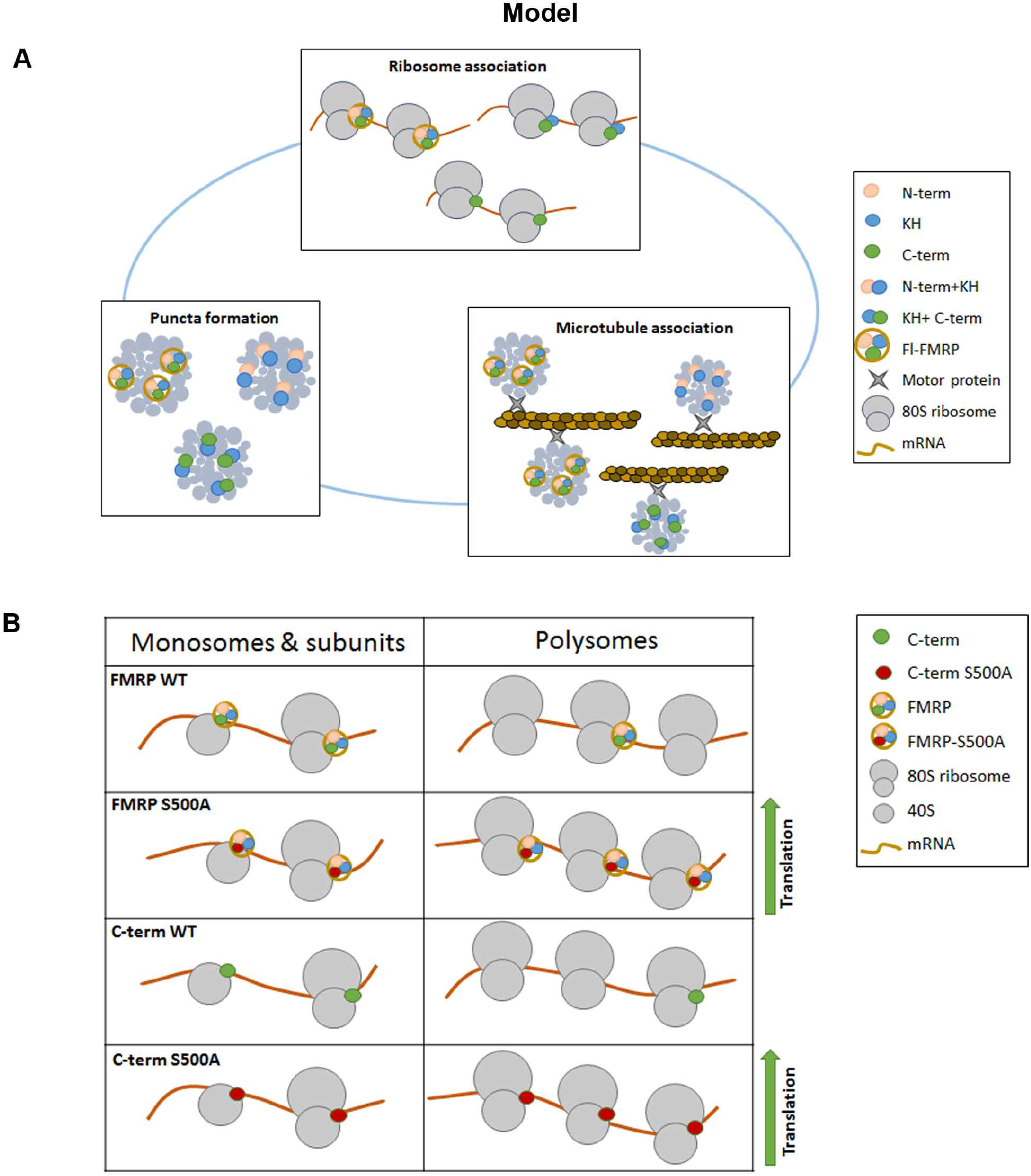
A. Model summarizing the minimal domains of FMRP required for puncta formation, microtubule association and association with ribosomes. B. Model indicating that the C-terminus is sufficient to regulate the translation function of FMRP in neurons. This function is further regulated through the dephosphosphorylation of FMRP, which increases translation through increased association with polysomes.

Our primary observation is that FMRP co-fractionates with ribosomal as well as non-ribosomal fractions (including mRNPs). This direct binding to the ribosome blocks the binding of essential translational machinery, facilitating the inhibition of translation (37). Our results describe an overall inhibition of protein synthesis in neurons expressing full-length FMRP. Independent studies separately claim that the KH and C-terminus domains are involved in this FMRP-mediated translation inhibition (24,37) but our experimental observations support the C-terminus in translational inhibition, analogous to the full-length FMRP (**Fig 7B**).

It is noteworthy that FMRP exists as a mixture of both phosphorylated and dephosphorylated forms in cells and phosphorylation is shown to influence the translation state of FMRP-associated ribosomes /polysomes (18,19). Through polysome profiling, we observe that phosphorylated FMRP gets enriched in the initial fractions of the gradient. Hence phosphorylation seems to redistribute FMRP into repressive granules. Stimulation of Gp1 mGluRs is shown to rapidly cause dephosphorylation of FMRP, a key feature in activity-mediated protein synthesis (18,19). Correspondingly, our study describes a key role for phosphorylation as a translational switch and how FMRP dephosphorylation (FMRP S500A) can activate protein synthesis by presumably increasing its ribosome association. Our findings also emphasize that a similar mechanism of translation regulation exists between dephosphorylated forms of FMRP and C-term, where both increase translation (FMRP S500A and C-term S500A) (**Fig 7B**). However, we could not capture the same molecular behavior between the phosphorylated forms of FMRP and C-term (FMRP S500D and C-term S500D).

In neurons, FMRP regulates dendritic mRNA translation to maintain multiple forms of plasticity (11,12). FMRP-bound target mRNA are shown to phase-separate into membrane-less granules in the cytoplasm and this is shown to be transported in a microtubule-dependent manner (38,39). The C-terminus has been extensively shown to drive FMRP granules/ puncta formation *in-vitro* (39). Despite these observations, the role of FMRP’s domains in granule formation and cytoskeleton-mediated transport is still unknown. Our data shows that the individual domains of FMRP are incapable of forming neuronal puncta on their own. Correspondingly, neither of the individual domains could single-handedly contribute to FMRP-microtubule interaction. Formation of mRNP granules and their consequent dendritic transport is mediated through a combination of protein-protein as well as protein-RNA interactions. Hence we speculate that the combination of domains improves microtubule association as well as puncta formation in comparison to their respective single domains (**Fig 7A**).

Neuronal granules have been previously shown to contain components of translation machinery like ribosomes (14). However the translation status of ribosomes within these granules is unknown. As shown previously, it is evident that phosphorylation of FMRP induces inhibition of translation (**Fig 3F**). Also, phosphorylation is shown to increase FMRP’s phase-separation propensity while dephosphorylation favors granule disassembly (39). We show that FMRP exists in phosphorylated and dephosphorylated states within granules and the area of the granules are dictated by the extent of phosphorylation. Additionally, our initial results postulate that phosphorylation might also influence the extent of ribosome binding within these granules. Inhibition of translation with Puromycin showed a striking reduction in the area and number of FMRP-containing granules and an even larger reduction with granules containing dephosphorylated FMRP. This suggests that a majority of granules containing FMRP also contain ribosomes, which are in an active state of translation elongation. FMRP-mediated local protein synthesis dictates that selected mRNA are stored in a repressed state with stalled polysomes within granules (40). Following an appropriate signal, this repression is immediately relieved to provide the bulk of locally required proteins. Our data offers support to a contrasting hypothesis whereby stored messages can also be actively translated when accumulated within granules (41).

The canonical mutation leading to FXS is the CGG-repeat expansion in the 5’UTR of the *FMR1* gene. But several sequence variants of FMR1 have also been shown to cause FXS-like symptoms (28–30,42,43). While most of these mutations have pathophysiological consequences, the molecular functions affected by them are unexplored. Through the use of point mutations, our study describes a hierarchical role of domains in cellular pathways that govern translation. Our results show that the C-terminus domain is important for ribosome association and mutations in the C-terminus domain (*G482S* and *R534H*) generate loss-of-function FMRP variants incapable of inhibiting translation. The other C-terminus frame-shift mutation we investigated *G538fs*23* depicted increased translation, which may be due to the altered amino acid sequence. FMRP with N-terminus (*R138Q*) and KH domain mutations (*I241N, 1304N* and *G266E*) also resulted in loss-of-function variants with respect to translation, although their C-terminus domain was unaffected. While the N-terminus and KH domain mutants failed to inhibit translation, they conversely increased global protein synthesis. This alternate mechanism of translation regulation adopted by the N-terminus and KH mutants is yet to be explored. The N terminus (*R138Q*) and KH mutant (*G266E*) proteins also fail to associate with microtubules, a function which likely precedes ribosome binding. Microtubule association comprises both RNA and protein interactions and the mutations could disrupt these properties, thus abolishing microtubule binding (27). In conclusion, distinct domains of FMRP contribute to independent roles in translation regulation. The mutations in the domains account for its inadequate performance in specific molecular functions, thus indicating a strong hierarchical role of FMRP domains.

## Materials and methods

### Ethics statement

All animal work was done in compliance with procedures approved by the Institutional Animal Ethics Committee (IAEC) and the Institutional Biosafety Committee (IBSC), InStem, Bangalore, India. Sprague Dawley (SD) rats were used in all animal experiments. Rodents were maintained at conditions with 20-22°C temperatures, 50-60 relative humidity, 0.3µm HEPA-filtered air supply at 15-20 ACPH and 14h/10h light/ dark cycle maintained.

### Cell line and primary neuronal cultures

HEK293T cells were cultured in Dulbecco’s Modified Eagle Medium (#21331–020 Thermo) containing 10% Fetal Bovine Serum (#F2442 Sigma) at 37°C and 5% CO_2_. Cells were passaged with 0.25% Trypsin solution (#15090046 Thermo) for 2 min and neutralized using the same culture medium.

Primary neuronal cultures were obtained from E18 rat embryos. Primary cultures were procedure was followed as previously described (8). Briefly, cortices from both the hemispheres were dissected out from E18 embryos in ice cold Hank’s Balanced Sat Solution (#H6648-1L Sigma). The tissue was incubated with 0.25% Trypsin solution for 5min at 37°C to enzymatically dissociate the cells and finally manually triturated in Minimum Eagles Medium (#<u>11095080</u> Thermo) containing 10% FBS (#F2442 Sigma) to generate a single cell suspension. Cells were plated at a density of 6×10^4^ cells / cm^2^ on glass coverslips coated with 0.2mg/ml Poly-L-lysine (#P2636-100MG Sigma) made in Borate buffer pH 8.5. Neurons were cultured for in MEM medium for 4h after plating following which they were cultured in Neurobasal Medium (#21103-049 Thermo) containing B27 Supplement (#17504-044 Thermo) and 1x Glutamax (#35050061 Thermo) for 11days at 37°C and 5% CO_2._ Cloning and Site Directed Mutagenesis

In plasmids generated for cell line and neuronal transfection, all truncations and site directed mutations (SDM) of FMRP were performed in the original plasmid pFRT-TODestFLAGHAhFMRPiso1 (Addgene #48690) containing full-length human FMR1 gene with N-terminal Flag-HA tag. Site directed mutagenesis was performed using overlapping PCR with Phusion® High-Fidelity DNA Polymerase (NEB M0530S) and primer sequences mentioned in **Table 1**. Plasmids generated for recombinant protein were constructed as follows. Sequences corresponding to N-terminus (1-216aa), KH (217-425aa) and C-terminus (425-632aa) were PCR amplified from plasmid pFRT-TODestFLAGHAhFMRPiso1 and cloned into PGEX6P1 vector (Cytiva # GE28-9546-48) with N-terminus GST tag and PreScission protease site. Full length FMR1 sequence was PCR amplified along with N-terminal 6XHis tag and cloned into PFastBac-HtB (Thermo #10712024) for Baculoviral protein expression.

**Table 1:**
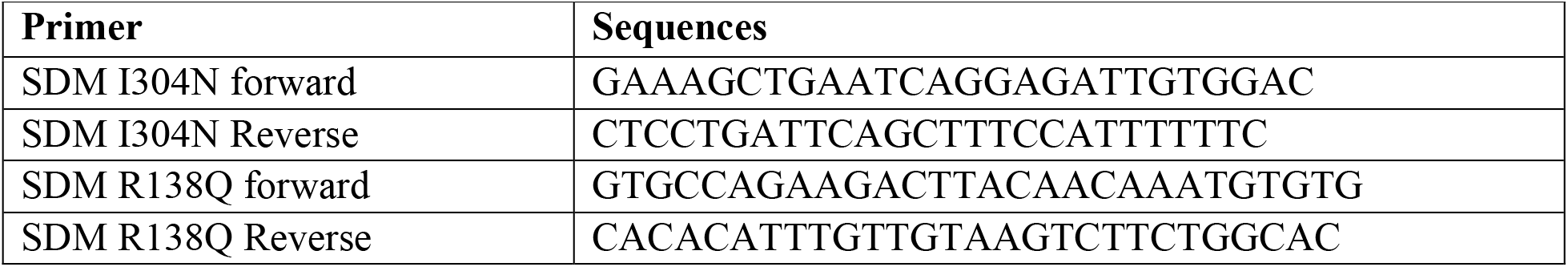
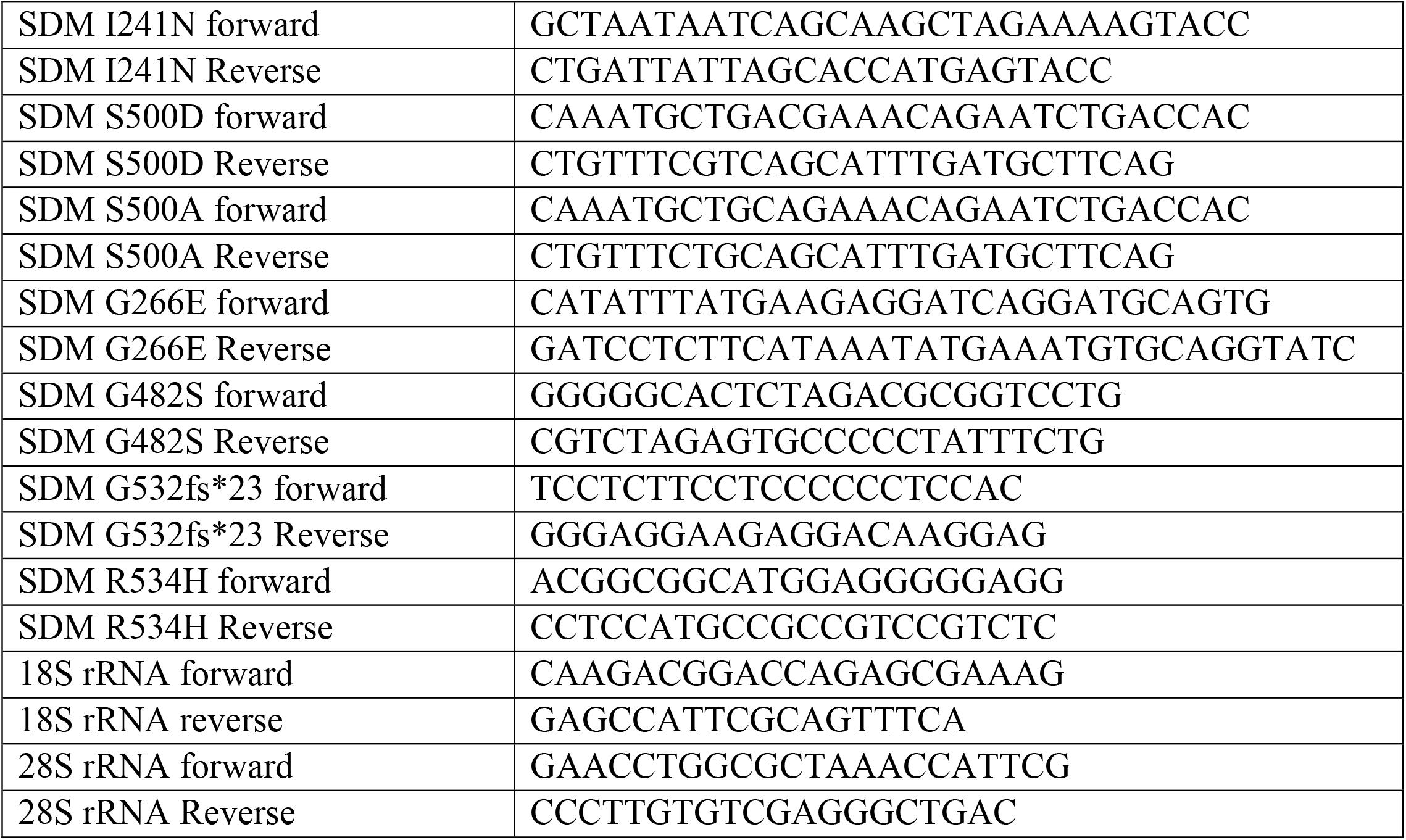
Primer details for SDM and qPCR

### Overexpression and protein purification experiments

For transfections, HEK293T cells/ DIV11 primary neurons were transfected with Flag-HA tagged FMRP constructs using Lipofectamine 2000(Thermo #11668027) for 24h as per the reagent’s protocol. His-GFP was also used as a control and transfection was performed in a similar manner as mentioned before. All constructs were transfected at equal concentrations. Flag-HA FMRP G538fs*23 showed very less transfection efficiency and hence was doubled in concentration with every transfection.

The N-terminus (1-216aa), KH (217-425aa) and C-terminus (425-632aa) of FMRP bearing GST tag at the N-terminal was transformed into *Escherichia coli* (*E*.*coli*) Rosetta DE3 cells and grown at 37 °C in LB. Protein expression was induced with 0.5 mM IPTG at an OD600nm ∼0.6 and grown overnight at 25 °C. Cells were harvested and pelleted via centrifugation. Pellets were stored at -20 °C or used immediately. Cells were lysed in buffer containing 20 mM Tris pH 8, 500 mM KCl, and 1mM DTT and sonicated for 30 min (10s ON, 10s OFF). Lysed cells were spun down and lysate was then loaded onto Glutathione Sepharose beads equilibrated with lysis buffer. Resin was washed with 10 column volumes of lysis buffer followed by 5 column volumes of lysis buffer with 100mM KCl. Elution buffer containing 50 mM Tris pH 8, 100mM KCl, 1mM DTT, 10% glycerol and 10mM reduced glutathione was used to elute the GST-tagged domains of FMRP. The flow-through was concentrated and further purified using a Superdex 75 column (GE Healthcare) equilibrated with buffer containing 50 mM Tris pH 8, 100mM KCl, 1mM DTT and 10% glycerol. Protein containing fractions were pooled together. The purity of protein was confirmed with SDS-PAGE gel.

Full length FMRP was cloned into Baculoviral pFastHTb vector with His-tag located towards the N-terminus of the protein. Sf9 insect cells were transfected with the construct with PEI and maintained at 26 °C at 120 rpm in SF900II medium (Thermo#10902088) until virus supernatant P0 was harvested 5 days post-transfection. Cell number, viability and diameter of the cells were monitored for Baculoviral infection. P1 virus supernatant was further harvested from a larger scale culture. P1 virus was used to infect a fresh culture of Sf21 cells. Baculoviral infection was monitored by recording cell number, viability and diameter prior to harvesting of cells. Cells were lysed in buffer containing 20mM Tris pH7.5, 800mM KCl, 5mM BME, 10%Glycerol and 10mM Imidazole and sonicated for 1min (10s ON, 10sOFF). Lysed cells were spun down and lysate was loaded onto Ni-NTA beads equilibrated with lysis buffer. Resin was washed with 10 column volumes of lysis buffer followed by 5 column volumes of lysis buffer. His-FMRP was eluted in buffer containing 20mM Tris pH7.5, 100KCl, 5mM β-ME, 10% Glycerol and 100mM Imidazole. The flow through was further dialysed in the same buffer to eliminate imidazole.

### Metabolic labelling

For metabolic labelling of transfected neurons, neurons were incubated in methionine-free Dulbecco’s Modified Essential Medium (Thermo# 21013024) for 30min followed by addition of azidohomoalanine (AHA; 1µM Thermo# C10102) in the same medium. This was incubated for 30min further and later fixed with 4%PFA. Cells were then permeabilised in PBS+0.3% Triton X-100 solution and blocked with buffer containing PBS+0.1% Triton X-100 + 2%BSA + 4% FBS solution. Newly synthesized proteins were then labelled with Alexa-Fluor-555–alkyne [Alexa Fluor 555 5-carboxamido-(propargyl), bis (triethylammonium salt)], by allowing the fluorophore alkyne to react with AHA azide group through click chemistry. All reagents were from Thermo Fisher, and stoichiometry of reagents was calculated according to the manufacture manual (CLICK-iT cell reaction buffer kit, #C10269). The neurons were subjected to immunostaining for MAP2B to identify neurons and HA to identify FMRP transfections (Antibody details in **Table 3**). The coverslips were mounted with Mowiol® 4-88 mounting media (#81381 Sigma) and imaged on Olympus FV3000 confocal laser scanning inverted microscope with 60X objective. The pinhole was kept at 1 Airy Unit and the optical zoom at 2X to satisfy Nyquist’s sampling criteria in XY direction. The objective was moved in Z-direction with a step size of 1 µM (∼8-10 Z-slices) to collect light from the planes above and below the focal plane. The transfected neurons were identified using the HA channel. Images of transfected and untransfected neurons were taken from each coverslip of every biological replicate. The image analysis was performed using ImageJ software and the maximum intensity projection of the slices was used for quantification of the mean fluorescent intensities. The region of interest (ROI) was drawn around the transfected and untransfected neurons using the MAP2 channel for their respective analyses. The mean fluorescent intensity of the FUNCAT channel was normalized to the MAP2 channel for each ROI. For each construct, the FUNCAT/MAP2 intensity ratio of the transfected neurons was normalized to the untransfected neurons from the corresponding biological replicate.

### Linear sucrose density centrifugation

Polysome assay was done from HEK293T cell lysate as described previously in (9). In brief, HEK293T cell lysate was separated on 15%–45% linear sucrose gradient in presence of 0.1mg/ml Cycloheximide (CHX) and Phosphatase inhibitor (Roche# 4906837001) by centrifugation at 39,000 rpm in SW41 rotor for 90 min. The sample was fractionated into 12 1.0 mL fractions with continuous UV absorbance measurement (A254). Fractions were further analysed by western blots. For quantification of FMRP distribution, Fractions were pooled (F3 and F4–11) according to subunit distribution based on RPLP0 and RPS6 immunoblotting. Similar protocol was followed with purified recombinant full length and domains of FMRP. 150pmoles of recombinant protein was spiked into pre-cleared HEK293T cell lysate and incubated for 10min on ice prior to separation on sucrose gradient.

### Binding assays

50pmoles of recombinant GST-tagged domains of FMRP was incubated with pre-cleared HEK293T cell lysate. Ribosome protein complexes were formed for 30min on ice and later incubated with 10µl of GST-Sepharose beads for 2h at 4°C. After incubation, beads were pelleted (30s at 1000 r.p.m.). The beads were then washed twice with 50 µl of ice-cold binding buffer (20mM HEPES pH7.5, 1mM DTT, 100mM KCl, 2.5mM MgCl_2_ and 10% glycerol). The amount of ribosome in cell lysate bound to beads was eluted with 1X Lamelli buffer and checked on a western blot. Ribosomes from HEK293T cells were purified as described in Khatter et al (44). 5pmoles of purified 80S was incubated with 50 pmoles of recombinant protein in 50µl of Binding buffer for 30min on ice. Ribosome-protein complexes were incubated with 10µl of GST-Sepharose beads for 2h at 4°C, eluted and quantified as mentioned previously. For quantification of rRNA, Trizol (Thermo#15596018) was added to the beads after the last wash and RNA was eluted as per manufacturer’s protocol. Total RNA was reverse transcribed into cDNA using MMLV Reverse transcriptase enzyme and random hexamers.

### Quantitative PCR

rRNA was quantified using specific primers (**Table 1**) designed against human 18SrRNA and 28SrRNA. Arbitrary copy numbers were calculated from a standard curve drawn from Ct values obtained from serial dilutions of cDNA for each candidate

### Microtubule enrichment assay

To separate microtubule polymers from free tubulin, HEK293T cells were treated with 10 nM paclitaxel (Sigma# PHL89806-10MG) for 1 h to stabilize microtubules. Cells were later lysed in buffer containing 150 mM KCl, 2 mM MgCl2, 50 mM Tris, pH 7.5, 2 mM EGTA, 2%glycerol, 10 nM paclitaxel, 0.125% Triton X-100, protease inhibitor cocktail (Sigma#S8830-20TAB), and 10 U of RNase-out (Thermo# 10777019) at room temperature for 5 min. To analyse microtubule association of the Flag HA-tagged wild type and mutant FMRP variants, HEK293T cells were transfected with these constructs 24h prior to paclitaxel treatment. For RNase treatment, the lysate buffer contained 1.2 µg/µl RNase A1 and 30 units of RNase T1 but no RNase-out. Nuclei were pelleted at 700 g for 1 min at room temperature, and the cytoplasmic supernatant was centrifuged for 10 min at 16,000 g at room temperature to pellet microtubule polymers. The microtubule pellet and the post microtubule supernatant were denatured in 1X Laemmli buffer followed by SDS-PAGE analysis. For microtubule-destabilizing experiments, HEK293Tcells were incubated with 2 µg/ml Nocodazole (Merck# M1404) for 15min prior to lysis with the same buffer containing Nocodazole and no paclitaxel.

### Western blotting

The elutes from in-vitro binding pull-downs, fractions of polysome profiling experiments and fractions from microtubule enrichment assay were analysed through western blotting. Briefly, the denatured lysates were run on 12% resolving and 5% stacking acrylamide gels and subjected to overnight transfer onto PVDF membrane. The blots were subjected to blocking for 1 hour at room temperature using 5% BSA prepared in TBST (TBS with 0.1% Tween-20). This was followed by primary antibody (prepared in blocking buffer) incubation at 4°C overnight. HRP tagged secondary antibodies were used for primary antibody detection. The secondary antibodies (prepared in blocking buffer) were incubated with the blots for 1 hour at room temperature. Three washes of 5-10 minutes each were given after primary and secondary antibody incubation using TBST solution. Details of primary and secondary antibody dilutions are provided in **Table 2**. The blots were subjected to chemiluminescent-based detection of the HRP tagged proteins. For the analysis of microtubule enrichment, the samples were probed with FMRP/GFP/HA antibody followed by α-Tubulin as the loading control. In cases where the over-expressed protein shared the same molecular weight as α-Tubulin, the blots were probed with FMRP/GFP/HA antibody first. The same blots were stripped and re-probed with α-Tubulin antibody later. For sucrose gradient fractions, distribution of overexpressed protein and RPLP0/RPS6 were analysed from the same blot. The details of the antibodies used and their dilutions are given in **Table 2**. All the western blot quantifications were performed using densitometry analysis on ImageJ software.

**Table 2:** Antibodies for Western Blot

| Antibody | Dilution | Catalog.no |
| --- | --- | --- |
| GST | 1:1000 | Ab9085, Abcam |
| 6His-Tag | 1:3000 | SAB4301134, Sigma |
| HA | 1:1000 | 3724S, CST |
| RPS6 | 1:3000 | 2217S, CST |
| RPLP0 | 1:3000 | Ab101279, Abcam |
| FMRP | 1:1000 | F4055, Sigma |
| p-FMRP | 1:1000 | Ab183319, Abcam |
| α-Tubulin | 1:3000 | T9026, Sigma |
| Secondary Rabbit HRP | 1:5000 | A0545, Sigma |
| Secondary Mouse HRP | 1:5000 | 31430, Thermofisher Scientific |

### Puncta analysis

Rat primary cortical neurons were transfected with HA-FMRP constructs on DIV11 followed by fixation with 4% PFA on DIV12. The fixed neurons were subjected to immunostaining for MAP2 and HA (Antibody details in **Table 3**). The coverslips were mounted with Mowiol® 4-88 mounting media (Sigma #81381) and imaged on Olympus FV300 confocal laser scanning inverted microscope with 60X objective. The pinhole was kept at 1 Airy Unit and the optical zoom at 2X to satisfy Nyquist’s sampling criteria in XY direction. The objective was moved in Z-direction with a step size of 0.5 µM (∼8-10 Z-slices) to collect light from the planes above and below the focal plane. The images were analysed using ImageJ software and the maximum intensity projection of the slices was used for quantification. The MAP2 channel was used to draw the ROI around each dendritic segment and subjected to thresholding. The ROI created using the MAP2 channel was selected on the HA-FMRP image. The area the selected ROI in the HA-FMRP channel was measured to obtain the total area of the dendritic ROI (a). Further, the ROI was subjected to particle analysis in the HA-FMRP image. Particles in the range of 3-30 pixels were considered for the analysis. The sum of the area of all the particles or the puncta area (b) and the number of particles or puncta (c) were the two parameters obtained using particle analysis. The ratio of puncta area (b) to total area (a) was calculated to measure the area occupied by the puncta in the given ROI. The ratio of puncta number (c) to total area (a) was calculated to measure the number of puncta per µm^2^. The ratios of puncta area/total area and number of puncta/ µm^2^ were compared among dendritic segment ROIs expressing different FMRP constructs.

**Table 3:**
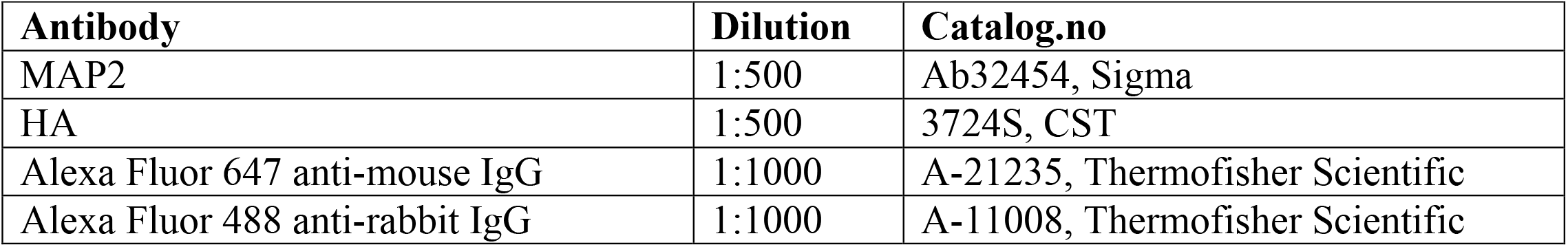
Antibodies for Immunostaining

| Antibody | Dilution | Catalog.no |
| --- | --- | --- |
| MAP2 | 1:500 | Ab32454, Sigma |
| HA | 1:500 | 3724S, CST |
| Alexa Fluor 647 anti-mouse IgG | 1:1000 | A-21235, Thermofisher Scientific |
| Alexa Fluor 488 anti-rabbit IgG | 1:1000 | A-11008, Thermofisher Scientific |

### Statistical analysis

All statistical analyses were performed using Graph Pad Prism software. The normality of the data was tested using the Kolmogorov-Smirnov test. For experiments with less than 5 data points, parametric statistical tests were applied. Data were represented as mean ± SEM in all in-vitro and polysome experiment graphs. FUNCAT and puncta data was represented as boxes and whiskers with all the individual data points. Statistical significance was calculated using Unpaired Student’s t-test (2 tailed with equal variance) in cases where 2 groups were being compared. One-way ANOVA was used for multiple group comparisons, followed by Tukey’s multiple comparison tests, Bonferroni’s multiple comparison test or Dunnett’s multiple comparison test. P-value less than 0.05 was considered to be statistically significant

## List of abbreviations

FMRP: Fragile X Mental Retardation Protein
RBP: RNA Binding Protein
RPS6: Ribosomal protein S6
RPLP0: Ribosomal Protein P0
NMDAR: *N*-methyl-D-aspartate receptor
mGLUR: Metabotropic glutamate receptor
FXS: Fragile X Syndrome
SNP: Single Nucleotide Polymorphism
FUNCAT: Fluorescent non-canonical amino-acid tagging
NLS: Nuclear Localization Signal
DMSO: Di-methyl Sulphoxide
ANOVA: Analysis of variance
E.coli: Escherichia coli
rRNA: Ribosomal RNA
mRNA: Messenger RNA
KH: K-Homology
WT: Wild type
UTR: Untranslated region
IPTG: Isopropyl ß-d-1-thiogalactopyranoside
BME: Beta-Mercaptoethanol
PBS: Phosphate Buffer Saline
MMLV: Moloney Murine Leukemia Virus
ROI: Region of interest
DTT: Dithiotheritol

## Author contributions

M.N.D., S.Ramakrishna, S.Ravindran and R.S.M.- designed the research; M.N.D., S.Ramakrishna, B.K.R and V.J.- performed the research; L.Y- contributed to experimental tools; M.N.D. and S.Ramakrishna- analyzed data; D.P. and R.S.M- provided funding; M.N.D- wrote the paper and M.N.D, S.Ramakrishna, S.Ravindran, B.K.R, V.J., D.P. and R.S.M reviewed versions of the paper.

## Acknowledgments

We thank Dr. Vinothkumar Kutti Ragunath, NCBS, for his useful comments and suggestions on the manuscript; Dr. Vinothkumar and Hitendra Negi, NCBS, for their help in our protein purification experiments; Sudhriti Ghosh Dastidar,inStem for his insights on FMRP puncta imaging and analysis; Central Imaging and Flow Cytometry Facility (CIFF) and Animal House Facility, NCBS-inStem; Members of Muddashetty lab and Palakodeti lab for their invaluable suggestions and discussions.

## References

1. Santoro MR, Bray SM, Warren ST. Molecular Mechanisms of Fragile X Syndrome: A Twenty-Year Perspective. Annu Rev Pathol Mech Dis. 2012 Feb 28;7(1):219–45.

2. Myrick LK, Hashimoto H, Cheng X, Warren ST. Human FMRP contains an integral tandem Agenet (Tudor) and KH motif in the amino terminal domain. Human Molecular Genetics. 2015 Mar 15;24(6):1733–40.

3. Nelson DL, Orr HT, Warren ST. The Unstable Repeats—Three Evolving Faces of Neurological Disease. Neuron. 2013 Mar;77(5):825–43.

4. Alpatov R, Lesch BJ, Nakamoto-Kinoshita M, Blanco A, Chen S, Stützer A, et al. A Chromatin-Dependent Role of the Fragile X Mental Retardation Protein FMRP in the DNA Damage Response. Cell. 2014 May;157(4):869–81.

5. Antar LN. Metabotropic Glutamate Receptor Activation Regulates Fragile X Mental Retardation Protein and Fmr1 mRNA Localization Differentially in Dendrites and at Synapses. Journal of Neuroscience. 2004 Mar 17;24(11):2648–55.

6. Darnell JC, Van Driesche SJ, Zhang C, Hung KYS, Mele A, Fraser CE, et al. FMRP Stalls Ribosomal Translocation on mRNAs Linked to Synaptic Function and Autism. Cell. 2011 Jul;146(2):247–61.

7. D’Souza MN, Gowda NKC, Tiwari V, Babu RO, Anand P, Dastidar SG, et al. FMRP Interacts with C/D Box snoRNA in the Nucleus and Regulates Ribosomal RNA Methylation. iScience. 2018 Nov;9:399–411.

8. Kute PM, Ramakrishna S, Neelagandan N, Chattarji S, Muddashetty Ravi S. NMDAR mediated translation at the synapse is regulated by MOV10 and FMRP. Mol Brain. 2019 Dec;12(1):65.

9. Muddashetty RS, Kelic S, Gross C, Xu M, Bassell GJ. Dysregulated Metabotropic Glutamate Receptor-Dependent Translation of AMPA Receptor and Postsynaptic Density-95 mRNAs at Synapses in a Mouse Model of Fragile X Syndrome. Journal of Neuroscience. 2007 May 16;27(20):5338–48.

10. Pasciuto E, Bagni C. SnapShot: FMRP Interacting Proteins. Cell. 2014 Sep;159(1):218-218.e1.

11. Huber KM, Gallagher SM, Warren ST, Bear MF. Altered synaptic plasticity in a mouse model of fragile X mental retardation. Proceedings of the National Academy of Sciences. 2002 May 28;99(11):7746–50.

12. Sidorov MS, Auerbach BD, Bear MF. Fragile X mental retardation protein and synaptic plasticity. Mol Brain. 2013;6(1):15.

13. Bear MF, Huber KM, Warren ST. The mGluR theory of fragile X mental retardation. Trends in Neurosciences. 2004 Jul;27(7):370–7.

14. Antar LN, Dictenberg JB, Plociniak M, Afroz R, Bassell GJ. Localization of FMRP-associated mRNA granules and requirement of microtubules for activity-dependent trafficking in hippocampal neurons. Genes Brain Behav. 2005 Aug;4(6):350–9.

15. Darnell JC. Kissing complex RNAs mediate interaction between the Fragile-X mental retardation protein KH2 domain and brain polyribosomes. Genes & Development. 2005 Apr 15;19(8):903–18.

16. Feng Y, Absher D, Eberhart DE, Brown V, Malter HE, Warren ST. FMRP Associates with Polyribosomes as an mRNP, and the I304N Mutation of Severe Fragile X Syndrome Abolishes This Association. Molecular Cell. 1997 Dec;1(1):109–18.

17. Stefani G. Fragile X Mental Retardation Protein Is Associated with Translating Polyribosomes in Neuronal Cells. Journal of Neuroscience. 2004 Aug 18;24(33):7272–6.

18. Muddashetty RS, Nalavadi VC, Gross C, Yao X, Xing L, Laur O, et al. Reversible Inhibition of PSD-95 mRNA Translation by miR-125a, FMRP Phosphorylation, and mGluR Signaling. Molecular Cell. 2011 Jun;42(5):673–88.

19. Narayanan U, Nalavadi V, Nakamoto M, Thomas G, Ceman S, Bassell GJ, et al. S6K1 Phosphorylates and Regulates Fragile X Mental Retardation Protein (FMRP) with the Neuronal Protein Synthesis-dependent Mammalian Target of Rapamycin (mTOR) Signaling Cascade. Journal of Biological Chemistry. 2008 Jul;283(27):18478–82.

20. Ceman S, O’Donnell WT, Reed M, Patton S, Pohl J, Warren ST. Phosphorylation influences the translation state of FMRP-associated polyribosomes. Human Molecular Genetics. 2003 Dec 15;12(24):3295–305.

21. Nalavadi VC, Muddashetty RS, Gross C, Bassell GJ. Dephosphorylation-Induced Ubiquitination and Degradation of FMRP in Dendrites: A Role in Immediate Early mGluR-Stimulated Translation. Journal of Neuroscience. 2012 Feb 22;32(8):2582–7.

22. Ferron L, Novazzi CG, Pilch KS, Moreno C, Ramgoolam K, Dolphin AC. FMRP regulates presynaptic localization of neuronal voltage gated calcium channels. Neurobiology of Disease. 2020 May;138:104779.

23. Zhan X, Asmara H, Cheng N, Sahu G, Sanchez E, Zhang F-X, et al. FMRP(1–297)-tat restores ion channel and synaptic function in a model of Fragile X syndrome. Nat Commun. 2020 Dec;11(1):2755.

24. Athar YM, Joseph S. The Human Fragile X Mental Retardation Protein Inhibits the Elongation Step of Translation through Its RGG and C-Terminal Domains. Biochemistry. 2020 Oct 13;59(40):3813–22.

25. Phan AT, Kuryavyi V, Darnell JC, Serganov A, Majumdar A, Ilin S, et al. Structure-function studies of FMRP RGG peptide recognition of an RNA duplex-quadruplex junction. Nat Struct Mol Biol. 2011 Jul;18(7):796–804.

26. Laggerbauer B. Evidence that fragile X mental retardation protein is a negative regulator of translation. Human Molecular Genetics. 2001 Feb 1;10(4):329–38.

27. Tekcan A. In Silico Analysis of FMR1 Gene Missense SNPs. Cell Biochem Biophys. 2016 Jun;74(2):109–27.

28. Collins SC, Bray SM, Suhl JA, Cutler DJ, Coffee B, Zwick ME, et al. Identification of novel FMR1 variants by massively parallel sequencing in developmentally delayed males. Am J Med Genet. 2010 Oct;152A(10):2512–20.

29. Handt M, Epplen A, Hoffjan S, Mese K, Epplen JT, Dekomien G. Point mutation frequency in the FMR1 gene as revealed by fragile X syndrome screening. Molecular and Cellular Probes. 2014 Oct;28(5–6):279–83.

30. Okray Z, Esch CE, Van Esch H, Devriendt K, Claeys A, Yan J, et al. A novel fragile X syndrome mutation reveals a conserved role for the carboxy-terminus in FMRP localization and function. EMBO Mol Med. 2015 Apr;7(4):423–37.

31. Suvrathan A, Hoeffer CA, Wong H, Klann E, Chattarji S. Characterization and reversal of synaptic defects in the amygdala in a mouse model of fragile X syndrome. Proceedings of the National Academy of Sciences. 2010 Jun 22;107(25):11591–6.

32. Pfeiffer BE, Huber KM. Fragile X Mental Retardation Protein Induces Synapse Loss through Acute Postsynaptic Translational Regulation. Journal of Neuroscience. 2007 Mar 21;27(12):3120–30.

33. Wang H, Dictenberg JB, Ku L, Li W, Bassell GJ, Feng Y. Dynamic Association of the Fragile X Mental Retardation Protein as a Messenger Ribonucleoprotein between Microtubules and Polyribosomes. Wickens MP, editor. MBoC. 2008 Jan;19(1):105–14.

34. Verkerk AJMH, Pieretti M, Sutcliffe JS, Fu Y-H, Kuhl DPA, Pizzuti A, et al. Identification of a gene (FMR-1) containing a CGG repeat coincident with a breakpoint cluster region exhibiting length variation in fragile X syndrome. Cell. 1991 May;65(5):905–14.

35. Ascano M, Mukherjee N, Bandaru P, Miller JB, Nusbaum JD, Corcoran DL, et al. FMRP targets distinct mRNA sequence elements to regulate protein expression. Nature. 2012 Dec;492(7429):382–6.

36. Brown V, Jin P, Ceman S, Darnell JC, O’Donnell WT, Tenenbaum SA, et al. Microarray Identification of FMRP-Associated Brain mRNAs and Altered mRNA Translational Profiles in Fragile X Syndrome. Cell. 2001 Nov;107(4):477–87.

37. Chen E, Sharma MR, Shi X, Agrawal RK, Joseph S. Fragile X Mental Retardation Protein Regulates Translation by Binding Directly to the Ribosome. Molecular Cell. 2014 May;54(3):407–17.

38. De Diego Otero Y, Severijnen L-A, van Cappellen G, Schrier M, Oostra B, Willemsen R. Transport of Fragile X Mental Retardation Protein via Granules in Neurites of PC12 Cells. Mol Cell Biol. 2002 Dec;22(23):8332–41.

39. Tsang B, Arsenault J, Vernon RM, Lin H, Sonenberg N, Wang L-Y, et al. Phosphoregulated FMRP phase separation models activity-dependent translation through bidirectional control of mRNA granule formation. Proc Natl Acad Sci USA. 2019 Mar 5;116(10):4218–27.

40. Darnell JC, Van Driesche SJ, Zhang C, Hung KYS, Mele A, Fraser CE, et al. FMRP Stalls Ribosomal Translocation on mRNAs Linked to Synaptic Function and Autism. Cell. 2011 Jul;146(2):247–61.

41. Mateju D, Eichenberger B, Voigt F, Eglinger J, Roth G, Chao JA. Single-Molecule Imaging Reveals Translation of mRNAs Localized to Stress Granules. Cell. 2020 Dec;183(7):1801-1812.e13.

42. Bechara EG, Didiot MC, Melko M, Davidovic L, Bensaid M, Martin P, et al. A Novel Function for Fragile X Mental Retardation Protein in Translational Activation. Wickens M, editor. PLoS Biol. 2009 Jan 20;7(1):e1000016.

43. Myrick LK, Nakamoto-Kinoshita M, Lindor NM, Kirmani S, Cheng X, Warren ST. Fragile X syndrome due to a missense mutation. Eur J Hum Genet. 2014 Oct;22(10):1185–9.

44. Khatter H, Myasnikov AG, Natchiar SK, Klaholz BP. Structure of the human 80S ribosome. Nature. 2015 Apr;520(7549):640–5.

