## Supplementary figures and legends for "Function of FMRP domains in regulating distinct roles of neuronal protein synthesis"

### Supplementary figure 1

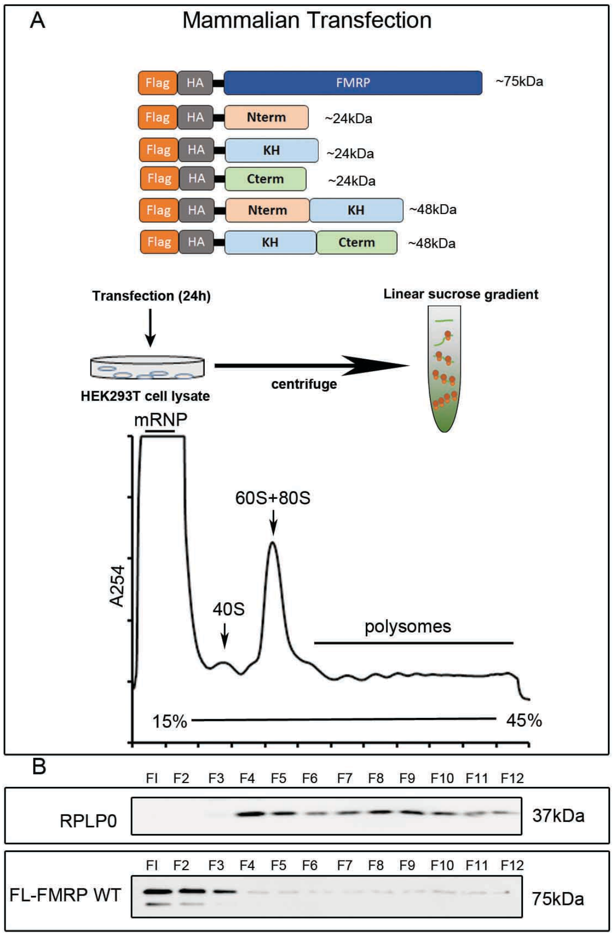


A. Schematic describing human FMRP constructs alongside truncated versions that were transfected in HEK293T cells followed by polysome profiling.
B. Representative immunoblots indicating the distribution of ribosomal protein RPLP0 (probed with anti-RPLP0 antibody) and overexpressed full-length FMRP (probed with anti-HA antibody) on linear sucrose gradient (n=3).

**Supplementary figure 2**

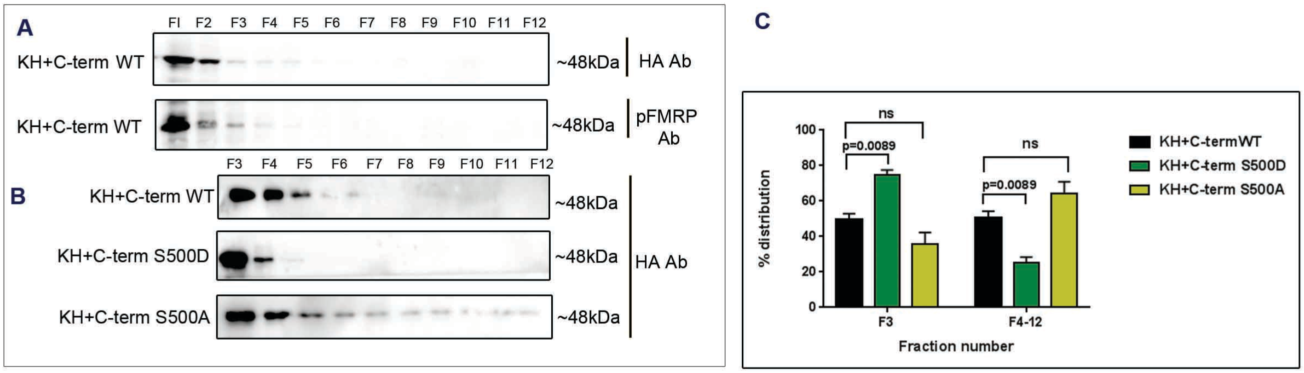


A. Representative immunoblots indicating the distribution of KH+C-term WT, phospho-mimetic (KH+Cterm S500D) and dephospho-mimetic (KH+Cterm S500A) overexpressed in HEK293T cells along all 12 fractions of linear sucrose gradient. Blot was probed with anti-HA antibody. Fractions were also probed for phospho KH+Cterm distribution using p-FMRP antibody (n=3).
B. Representative immunoblots indicating the distribution of overexpressed KH+Cterm WT and phospho-mutants along fractions 3 to 12 of linear sucrose gradient. Blots were probed with anti-HA antibody (n=3).
C. Graph indicating the quantification of overexpressed KH+Cterm variants in non-ribosomal (F3) versus Ribosomal fractions (F4-12). n=3 for each condition. Data represented as mean +/- SEM, One way ANOVA p=0.001 for F3, p=0.001 for F4-12, Followed by Bonferroni's Multiple comparisons test

### Supplementary figure 3

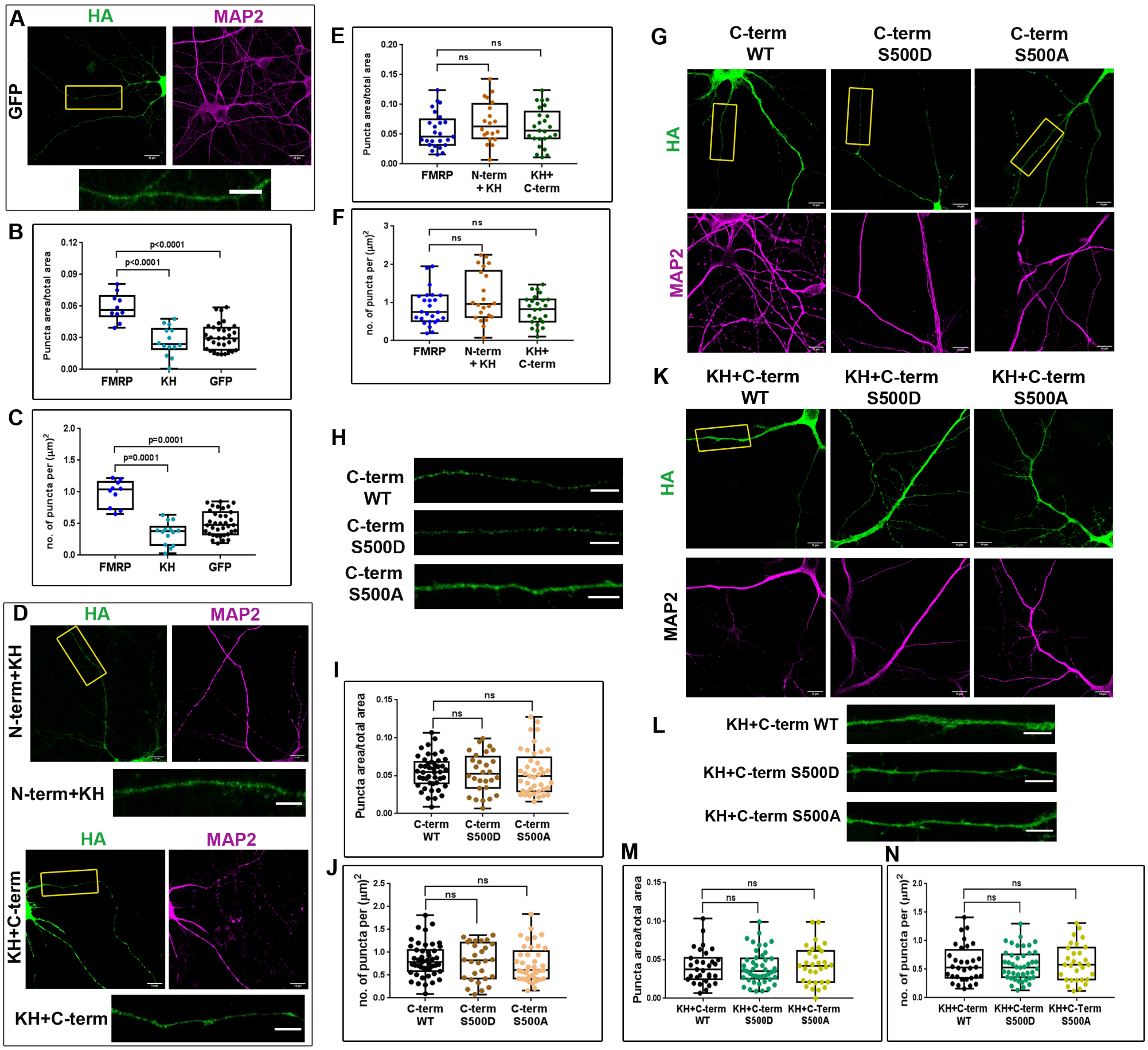


A. Representative images of HA and MAP2 fluorescent intensities in Primary rat cortical neurons (DIV11) transfected with control GFP, N-term+KH and KH+C-term for 24h (Scale bar - 10μm).
B. Inlets shown in panel A are enlarged showing dendritic localization of puncta (Scale bar - 5μm).
C. Box plot representing the quantification of puncta area by total dendritic area for length FMRP, GFP and KH domain of FMRP. The box extends from 25th to 75th percentile with the middlemost line representing the median of the dataset. Whiskers range from minimum to maximum data point. One way ANOVA $p<0.001$ followed

by Tukey’s multiple comparison’s test, n = 11-35 neurons from 3 independent experiments.

D. Box plot representing the quantification of the number of puncta per unit area of the dendrite for full length FMRP, GFP and KH domain of FMRP. The box extends from 25th to 75th percentile with the middlemost line representing the median of the dataset. Whiskers range from minimum to maximum data point. One way ANOVA $p<0.001$ followed by Tukey’s multiple comparison’s test, n = 11-35 neurons from 3 independent experiments.

E. Box plot representing the quantification of puncta area by total dendritic area for length full length FMRP, N-term+KH and KH+C-term of FMRP. The box extends from 25th to 75th percentile with the middlemost line representing the median of the dataset. Whiskers range from minimum to maximum data point. One-way ANOVA $p=0.3415$ followed by Tukey’s multiple comparison’s test, n = 20-25 neurons from 5 independent experiments.

F. Box plot representing the quantification of the number of puncta per unit area of the dendrite for full length FMRP, N-term+KH and KH+C-term of FMRP. The box extends from 25th to 75th percentile with the middlemost line representing the median of the dataset. Whiskers range from minimum to maximum data point. One-way ANOVA $p=0.0566$ followed by Tukey’s multiple comparison’s test, n= 20-25 neurons from 5 independent experiments.

G. Representative images of HA and MAP2 fluorescent intensities in Primary rat cortical neurons (DIV11) transfected with WT, S500D and S500A C-term variants for 24h (Scale bar - 10μm).

H. Inlets shown in panel G are enlarged showing dendritic localization of puncta (Scale bar - 5μm).

I. Box plot representing the quantification of puncta area by total dendritic area for full-length WT, S500D and S500A C-term. The box extends from 25th to 75th percentile with the middlemost line representing the median of the dataset. Whiskers range from minimum to maximum data point. n= 25-45 neurons from 5 independent experiments. One-way ANOVA $p=0.02238$ followed by Tukey’s multiple comparison test.

J. Box plot representing the quantification of the number of puncta per unit area of the dendrite for full length and WT, S500D and S500A C-term. The box extends from 25th to 75th percentile with the middlemost line representing the median of the dataset. Whiskers range from minimum to maximum data point. n= 25-45 neurons from 5 independent experiments. One-way ANOVA $p=0.5737$ followed by Tukey’s multiple comparisons test.

K. Representative images of HA and MAP2 fluorescent intensities in Primary rat cortical neurons (DIV11) transfected with WT, S500D and S500A KH+C-term variants for 24h (Scale bar - 10μm).

L. Inlets shown in panel K are enlarged showing dendritic localization of puncta. (Scale bar - 5μm).

M. Box plot representing the quantification of puncta area by total dendritic area for full length WT, S500D and S500A KH+C-term. The box extends from 25th to 75th percentile with the middlemost line representing the median of the dataset. Whiskers range from minimum to maximum data point. n= 30-45 neurons from 5 independent experiments. One-way ANOVA p=0.1518 followed by Tukey's multiple comparison test.

N. Box plot representing the quantification of the number of puncta per unit area of the dendrite for full length and WT, S500D and S500A KH+C-term. The box extends from 25th to 75th percentile with the middlemost line representing the median of the dataset. Whiskers range from minimum to maximum data point. n= 30-45 neurons from 5 independent experiments. One-way ANOVA p=0.199 followed by Tukey's multiple comparisons test.

**Supplementary figure 4**

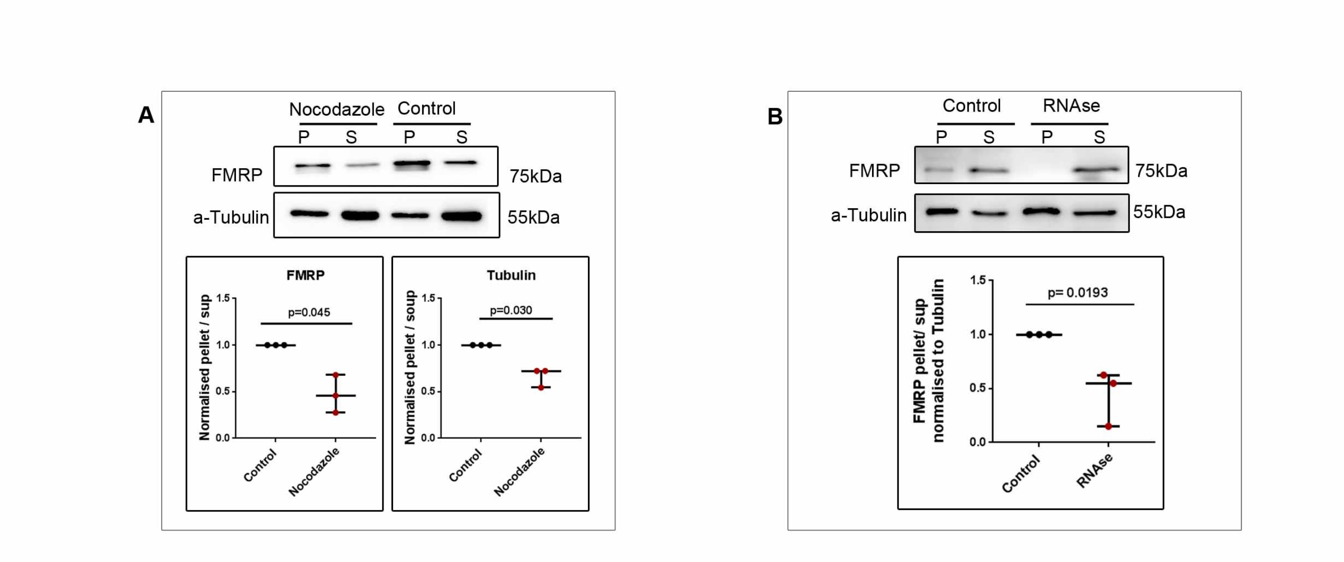


A. Top - Representative immunoblots indicating the de-enrichment of endogenous FMRP and Tubulin in microtubule pellet on Nocodazole treatment in HEK293T cells. Bottom - Graphs indicating the pellet to supernatant ratio of endogenous FMRP (left) and Tubulin (right) in Nocodazole versus DMSO (Control) treated cells. Data represented as mean +/- SEM, Unpaired t-test (n=3).

B. Top – Representative immunoblots indicating the de-enrichment of endogenous FMRP in microtubule pellet on RNase treatment in HEK293T cells. Bottom - Graph indicating the pellet to supernatant ratio of endogenous FMRP in RNase treated cells. Data represented as mean +/- SEM, Unpaired t-test (n=3).
